# Actin in Dendritic Spines Self-Organizes into a Critical State

**DOI:** 10.1101/2020.04.22.054577

**Authors:** Mayte Bonilla-Quintana, Florentin Wörgötter, Elisa D’Este, Christian Tetzlaff, Michael Fauth

## Abstract

It is known that dendritic spines change their size and shape spontaneously and sometimes to a large degree, but the function of this remains unclear. Here, we quantify these changes using time-series analysis of confocal data and demonstrate that spine size can follow different autoregressive integrated moving average (ARIMA) models and that shape- and size-changes are not correlated. We capture this behavior with a biophysical model, based on the spines’ actin dynamics, and find the presence of 1*/f* noise. When investigating its origins, the model predicts that actin in the dendritic spines self-organizes into a critical state, which creates a fine balance between static actin filaments and free monomers. We speculate that such a balance might be functionally beneficially to allow a spine to quickly reconfigure itself after LTP induction.

## Introduction

Dendritic spines are small protrusions of dendrites where the postsynaptic part of most excitatory synapses is located. It is well known that size and shape changes of these spines are correlated with changes of the strength of excitatory synaptic connections (Matsuzaki et al., 2004). However, Fischer et al. (1998) observed that the shape of dendritic spines also varies spontaneously during 30-minute recordings at a 1.5-second frame-rate under basal conditions, under which spines are at the minimum level of activity required to maintain their functions. Interestingly, nearly all spines in these recordings survived and their volume and density stayed mostly constant. Similar spine variations were measured by Dunaevsky et al. (1999) in time-lapse images of acute and cultured slices from cerebellar Purkinje cells, and cortical and hippocampal pyramidal cells. Rapid spine shape changes have also been observed in living organotypic hippocampal mouse brain slices (Testa et al., 2012) and in the somatosensoty cortex of the adult mouse in vivo (Berning et al., 2012).

Rapid changes of dendritic spine shapes are related to actin dynamics (Fischer et al., 1998; Dunaevsky et al., 1999). Actin is a globular protein that assembles into filaments, which are polar structures that continuously undergo a treadmilling process. In this process, actin monomers are polymerized at the (+) end (barbed end) of the actin filaments, while these filaments are depolymerized at the (-) end. Actin polymerization generates a force that moves the cell membrane forward (Mogilner & Oster, 1996). Moreover, blockage of actin polymerization hinders spine motility (Fischer et al., 1998; Dunaevsky et al., 1999). Honkura et al. (2008) observed filamentous actin within a single spine and found that a dynamic pool of actin, which has a fast treadmilling velocity and is mainly localized at the tip of the spine, can account for fast spine motility. However, Frost et al. (2010) observed that actin treadmilling velocity is elevated not only at the spine tip, but also in discrete and well-separated foci.

There are numerous theoretical and experimental studies of dendritic spine size changes. However, these studies mostly consider spine development (Hotulainen et al., 2009; Miermans et al., 2017), the effects of long-term potentiation (Fauth et al., 2015), or spine volume fluctuations but at longer time scale (Yasumatsu et al., 2008; Dvorkin & Ziv, 2016). Recently, Bonilla-Quintana et al. (2020) proposed an actin based model that mimics rapid spine motility. However, to date there is no general explanation for the possible function of these rapid shape fluctuations (Chazeau & Giannone, 2016).

Here, we study rapid, spontaneous shape fluctuations of dendritic spines in experimental time-lapse data previously published in (Mikhaylova et al., 2018) as well as in new data from the lab of Elisa D’Este (abbreviated in the following by E.D.). We quantify size and shape changes by employing in a new way a method from circular statistics and show that these characteristics can be captured by a biophysical model. This model describes the interaction of actin treadmilling with the spine’s membrane (Bonilla-Quintana et al., 2020) also assuming that there are slow dynamic processes that alter the structure of the PSD and the spine neck. The model allows us to predict spine characteristics for periods of time that exceed the currently accessible experimental durations. Here we find that actin polymerization occurs in bursts from specific polymerization foci and is, possibly, best described by a process of self-organized criticality (Bak et al., 1987) also observed in other neural substrates (Beggs & Plenz, 2003). Since spine size correlates to synaptic strength (Matsuzaki et al., 2004), the spine cytoskeleton must enlarge upon LTP induction. An actin polymerization process that operates in a critical state, as suggested by the current study, may be functionally beneficial for achieving such plasticity-induced spine restructuring most efficiently.

## Results

In order to study spine shape fluctuations, we re-analysed basal data from primary mouse neurons at 12 days in vitro (DIV12) previously published in (Mikhaylova et al., 2018) and kindly provided by Dr. Marina Mikhaylova. These neurons were transfected with mRuby2 as cell fill and GFP-actin as actin marker, and imaged every 10 seconds for 5 minutes. Using Fiji (Schindelin et al., 2012), we traced a region of interest (ROI) around the heads of 16 spines for each time frame (yellow polygon in Fig. 1A). We noted that spine shape varies substantially over time, even for consecutive time frames.

**Figure 1:**
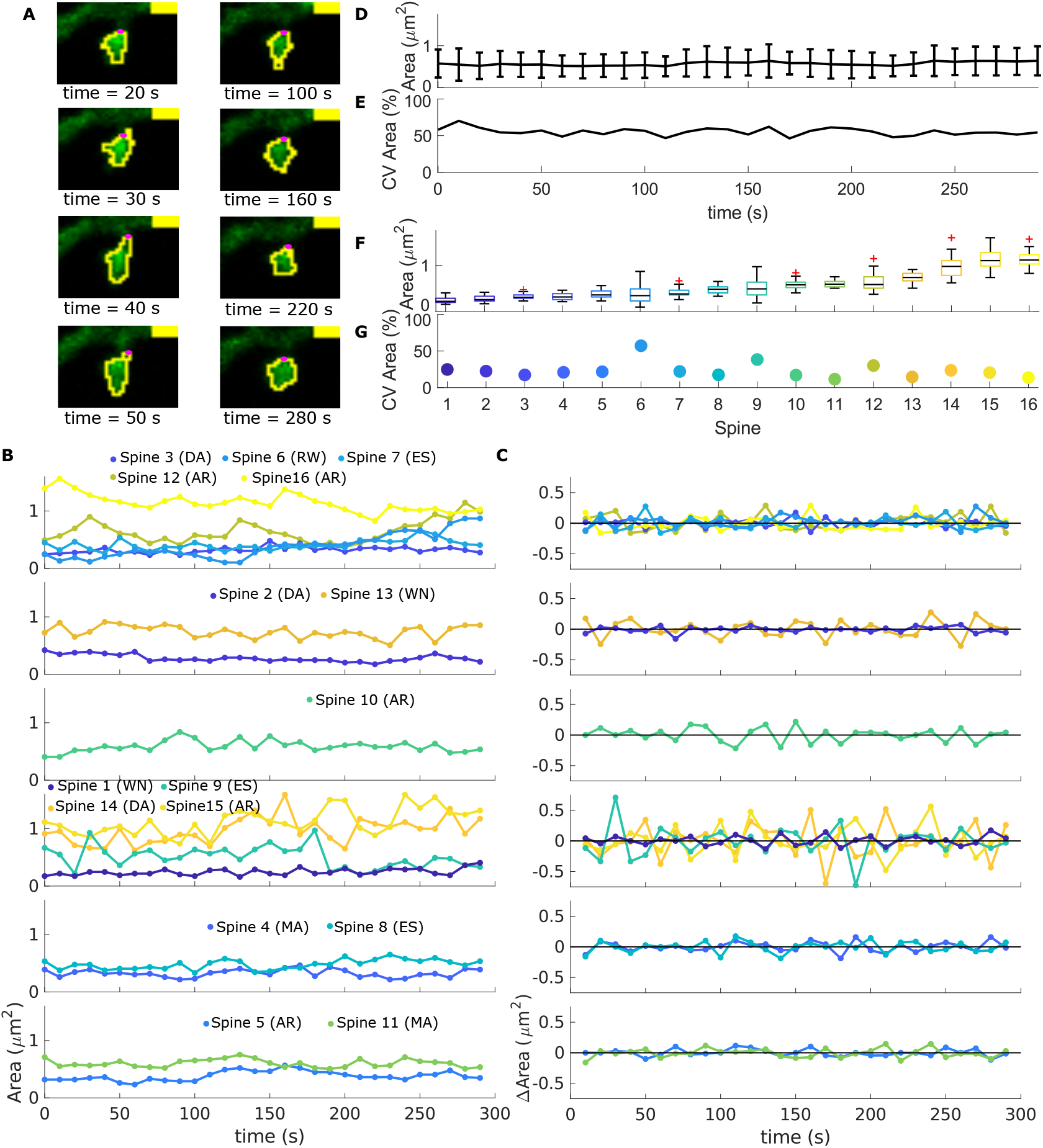
Spine evolution over time. **A** Images taken from time-lapse data of neurons transfected with GFP-actin (green). The yellow polygon represents the analysed ROI and the magenta circle corresponds to the spine neck center. Yellow bar indicates 1*μ*m. **B** Area evolution for spines in different stacks (top to bottom). Spines are color-coded and numbered from small (dark blue) to large (yellow). Abbreviations correspond to the best fit of an ARIMA model (see Table S1). **C** Difference of area, ΔArea, from spines in B over time. **D** Mean ± standard deviation (std) of the area of the analysed spines at each time-frame. **E** Coefficient of variation (CV) from D. **F** Boxplots of spine area evolution for each spine. **G** CV from F.

### Spine area evolution over time shows different tendencies

To investigate if spine size is preserved during the recordings regardless of shape changes, as observed Fischer et al. (1998), we analyzed the spine area evolution over time (Fig. 1B). In the same stack (i.e., image set of the same frame taken at different times), spines with different sizes and fluctuation behavior were observed. The average area of the spines spreads over a continuous range and the variability does not correlate to spine size (Fig. 1F). To quantitatively test the latter, we calculated the coefficient of variance (CV) for each spine time evolution (see Fig. 1G). In sum, we observed that spines behave differently and without any correlation to their size. For example, big spines exhibit large (Spine 12), but also small (Spine 16) area fluctuations that can be either fast (Spine 15) or slow (Spines 12, 16). And the same is true for small spines (compare Spine 6 with Spines 1, 5). However, the average (Fig. 1D) and variability (Fig. 1E) of the spine sizes at each frame were conserved.

When examining the spine area differences between successive time frames ΔArea(*t_i_*) = Area(*t_i_*) *−* Area(*t_i−_*_1_) (Fig. 1C), we noted a tendency to counteract a positive change in area by a negative change. Hence, the spine area tends to fluctuate around a mean value during the analyzed 5-minute recordings. To quantitatively describe this area tendency, we performed a time series analysis using autoregressive integrated moving average (ARIMA) models (see details in Methods). Thus, possible tendencies in area evolution express themselves through which ARIMA model best fits the data. We obtained that the area of 5 spines (31.25%), best fitted by an ARIMA(1,0,0), tends to return to a constant area mean value, either rapidly (Spines 10, 15) or slowly (Spines 5, 12, 16). Although this was the most common tendency, some spines show different behavior (see Table S1), which could also be due to the limited duration of the experiments or the lack of finer time resolution. Both constraints may result in a sub-optimal sampling of the spine area evolution and can lead to a poorer fitting of the ARIMA models.

### Spine shape changes are not correlated to size changes

Besides changes in spine size, we observed high variability in spine shapes. To determine whether there is a correlation between these fluctuations, we proposed shape descriptors that account for changes in the directional and orientational structure of the spine head using concepts from circular statistics (Batschelet, 1981). Such descriptors can detect if the spine elongates or tilts towards a certain direction. This can be used to quantify different shape configurations.

### Shape Descriptors

For the proposed shape descriptors, we took the spine neck center as the origin point and traced a ray every 15 degrees (*≈* 0.2618 rad). If the ray corresponding to the angle *θ_i_* intersected the ROI, then a sample point was taken at the intersection (black dot in Fig. 2A) and the distance to neck center was measured (dROI(*θ_i_*) brown line in Fig. 2B). If the traced ray did not intersect the ROI, then dROI(*θ_i_*) = 0. Thus, dROI(*θ*) accounts for the distance from the neck center to the spine membrane at the sample points. Note that dROI(*θ*) has a 2*π* period and each spine has the same number of sampled points regardless of size. To obtain an expression for dROI(*θ*), we performed optimal periodic regression (see Methods). Figure 2B shows the resulting function *R*(*θ*) that is also periodic, and hence, can be divided into three parts:

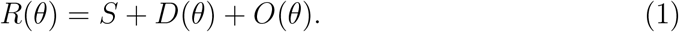

**Figure 2:**
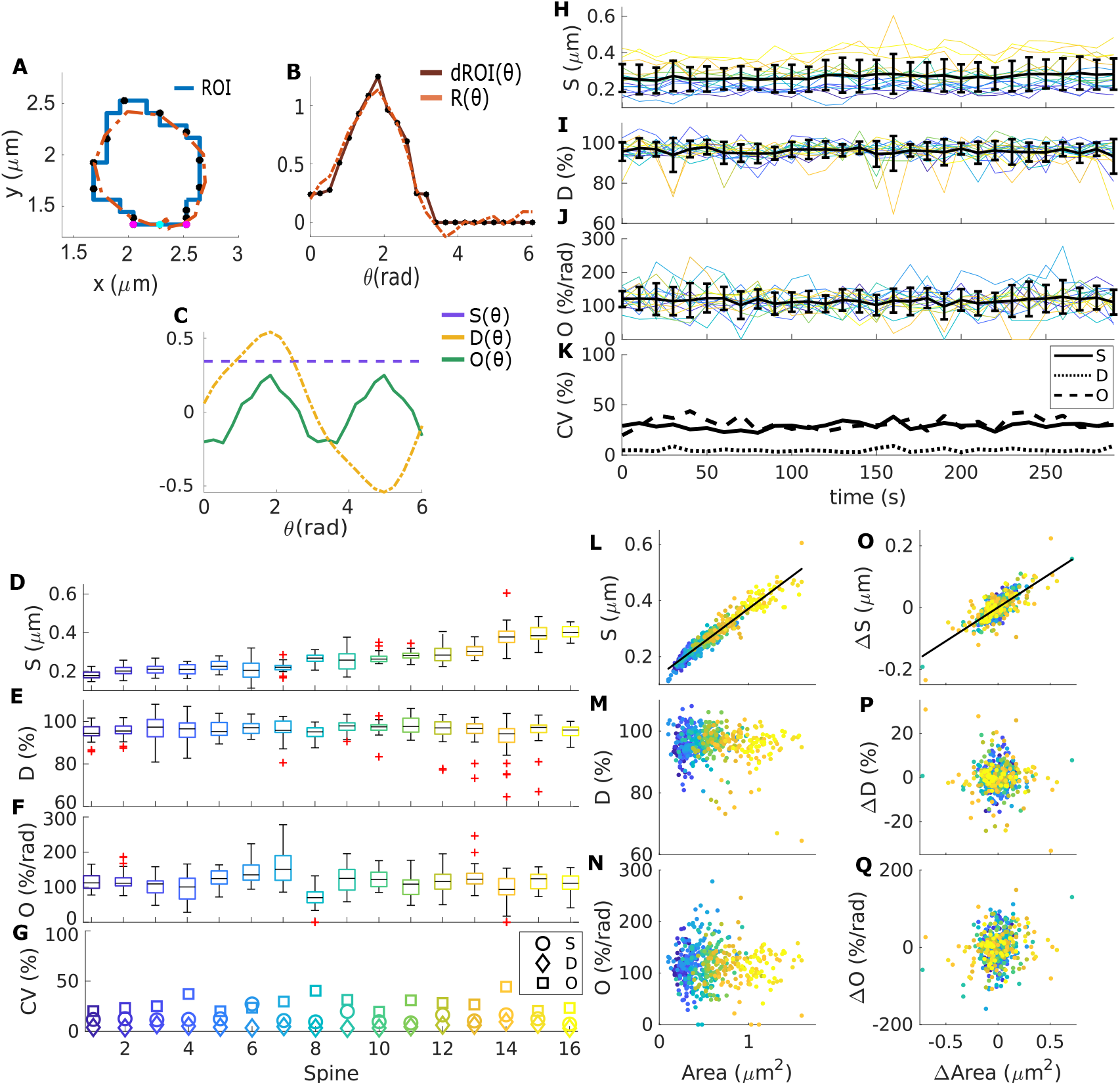
Shape descriptors. **A** Blue line: ROI extracted from data corresponding to the spine in Fig. 1A at time = 280 seconds. Black dots correspond to the sampled points of the ROI and the orange dotted line to the approximation function *R*(*θ*). Magenta dot denotes the spine neck center and cyan dots the neck limits. **B** Plot of the distance between the neck and sample points dROI(*θ*) (brown line) and *R*(*θ*) (orange line) over the sampling angles *θ_i_* (in radians). **C** The three parts of *R*(*θ*) corresponding to the general size of the spine *S*(*θ*), the directional selectivity *D*(*θ*), and orientational selectivity *O*(*θ*). **D-F** Box plot of shape descriptors for the analyzed spines, color-coded as in Figure 1. **G** Coefficient of variance of the shape descriptor for each spine. **H-J** Shape descriptors, *S*, *D*, and *O*, evolution over time for different spines. **K** Coefficient of variance corresponding to H-J. **L-N** Spine area against shape descriptors *S*, *D* and *O*, respectively. Each dot represents the corresponding value at a time-frame for a certain spine (color-coded). Black line is a linear least-squares fit. Only *S* correlates significantly with the area (p-value *<* 0.01; Pearson’s linear correlation coefficient cc= 0.96). **O-Q** Same as L-M but using the time differences of the shape descriptors. Δ*S* correlates with ΔArea (cc= 0.77).

These parts correspond to general size *S* of the spine, the preference of the spine to tilt along a certain direction *D*(*θ*) and the orientational selectivity of the spine head *O*(*θ*) (see Fig. 2 C). In order to describe spine shapes, *D*(*θ*) and *O*(*θ*) should be considered over the full period. Moreover, to compare different spines with different sizes, *D*(*θ*) and *O*(*θ*) should be calculated in terms of *S*. Thus, following Li et al. (1994), these quantities are integrated and averaged:

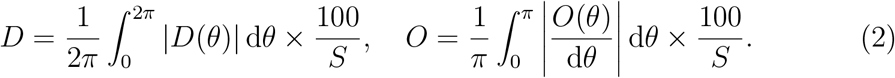

Note that *D* and *O* are expressed in terms of percentage of *S* and percentage of *S* per radian, respectively. These descriptors can thus characterize asymmetrical shape configurations. For example, *D* is bigger if the spine head considerably tilts towards one direction (from the spine neck), whilst *O* has larger values if the spine elongates along one or more directions. For further interpretation of these descriptors, see Supporting Information.

### Shape descriptors from experimental data

These shape descriptors were calculated for each spine at every time-frame. As shown in Figure 2H-J, the quantities fluctuate over time and, although each spine behaves differently (see Fig. 2D-G), they seem to fluctuate around a mean value with small changes in their variability (Fig. 2K). Moreover, the shape descriptors also seem to alternate between positive and negative changes; however, an analysis using ARIMA models shows that the fluctuations in orientational and directional selectivity mostly behave like white noise (50% of the spines), indicating that changes in *D* and *O* are not correlated in time. Further examination shows that only *S* significantly correlates to the spine area and that changes in area are only strongly correlated to those in *S* (Figs. 2L and O). Such correlations are expected since *S* accounts for the average distance from the spine neck to the membrane. Because *D* and *O* and their differences are not strongly correlated to the spine area and ΔArea, respectively (Figs. 2M-N and P-Q), we concluded that size fluctuations are independent to those in shape, in line with previous experimental results using different measures (Fischer et al., 1998).

To test whether these results are independent of the data set, we obtained and analyzed a new recording, consisting of 100 frames sampled every *≈* 9.6 seconds from hippocampal neurons DIV19 labelled with DiO provided by E.D. (see Methods). Figure 3A-D shows that this data behaves similarly: the evolution of the shape descriptors fluctuate around a mean value. Moreover, the area and its changes are only significantly correlated to *S* and its changes (cc= 0.5917 and cc= 39.49, with p-value *<* 0.01 using Student’s *t* distribution, respectively, see Figs. 3F and I). Note that the area values of the new data set correspond to a single spine in the previous data set (see Figs. 3F-K); hence, there is less variability in the area and *S*, but the variability in *D* and *O* is similar (Fig. 3E). Therefore, it appears that spines recorded with different experimental set-ups can behave similarly.

**Figure 3:**
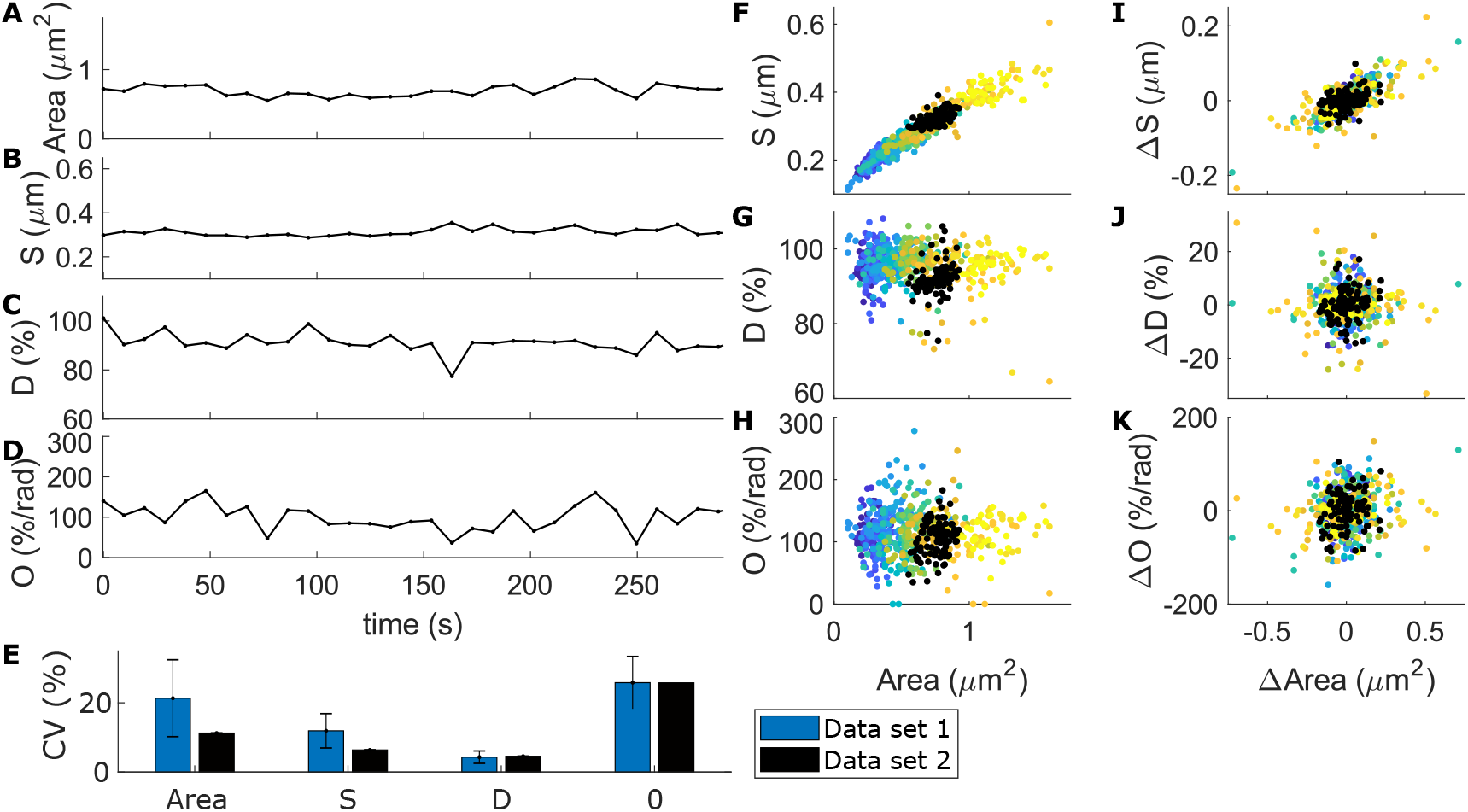
Comparison between different experimental data sets. Here the data set 1 corresponds to that of Figure 2, and data set 2 to a new one from the lab of E.D. **A-D** Area and shape descriptors (*S*, *D*, and *O*) evolution over time for a spine belonging to data set 1. **E** Median standard deviation of the coefficient of variability (CV) for shape descriptors calculated for each spine in data set 1 (blue) and CV for the spine from data set 2 (black). **F-H** Spine area against shape descriptors *S*, *D* and *O*. Each dot represent a measure of shape at a sampled time frame. There are 16 spines sampled every 10 seconds for 5 minutes corresponding to data set 1, color-coded as in Figure 2 and 100 frames from data set 2 sampled every 9.6 seconds (black). **I-K** Changes in area against changes in shape descriptors from the data in (F-H).

### Study of spine shape fluctuations using a theoretical model

So far, our study of dendritic spine shape fluctuations had been limited to the time and space resolution of the experiments. Thus, the different trends detected in the spine area and shape evolution could be due to the short duration or the limited number of frames in the recordings. To investigate whether a unique trend in the spine shape and size evolution can emerge over longer periods, we used a theoretical model that can be simulated for longer times and sampled as needed.

### Theoretical Model

We performed simulations based on the model from Bonilla-Quintana et al. (2020) (see Methods for model equations), in which asymmetric shape fluctuations emerge from an imbalance between a force generated by actin polymerization foci, that pushes the membrane forward at different locations, and a force generated by the membrane, that counteracts shape deformations.

This model considers the dynamics of actin filaments (F-actin) at the polymerization foci. Actin continuously undergoes a treadmilling process in which monomers (G-actin) are polymerized at the (+) end of the filaments (also called the “barbed end”), whilst G-actin is depolymerized or severed at the pointed (-) end (Fig. 4A). However, the ends of actin filaments can be capped when a capping protein binds to it, and thereupon, F-actin can not be polymerized (at the + end) or depolymerized (at the - end). When a focus of actin polymerization is initiated, it contains an actin filament with capped (-) end and uncapped (+) end. In addition to polymerizing G-actin at the barbed (uncapped +) end of a filament at each time-step, there are other events that can occur with a certain probability, namely, barbed ends can branch and form new filaments with barbed end and capped (-) end. This branching probability depends on the membrane force at that location and the number of barbed ends; hence it acts like a feedback mechanism that disables branching, that in turn lowers actin force, when the focus has pushed the membrane extensively or when there is an increased number of barbed ends. Additionally, barbed ends can be capped or capped minus ends can be uncapped at a constant rate, and filaments with uncapped (-) ends can be severed (see Fig. 4C). Therefore, the number of barbed ends at each focus fluctuates until they become extinct within seconds and the focus vanishes. To maintain fluctuations, new actin polymerization foci are nucleated with a given probability at a randomly chosen location near to the PSD (purple dot in Fig. 4B) that push the membrane in a chosen direction (thick arrow in Fig. 4B). The events of F-actin are modeled by a Monte Carlo scheme and the number of barbed ends is counted at each time-step to calculate the force generated by actin polymerization that is modeled by a Gaussian kernel proportional to them (see zoom in Fig. 4B). In addition to these basic model properties, we are also using mechanisms that allow slow and small displacements and changes of PSD as well as spine neck (see Methods).

**Figure 4:**
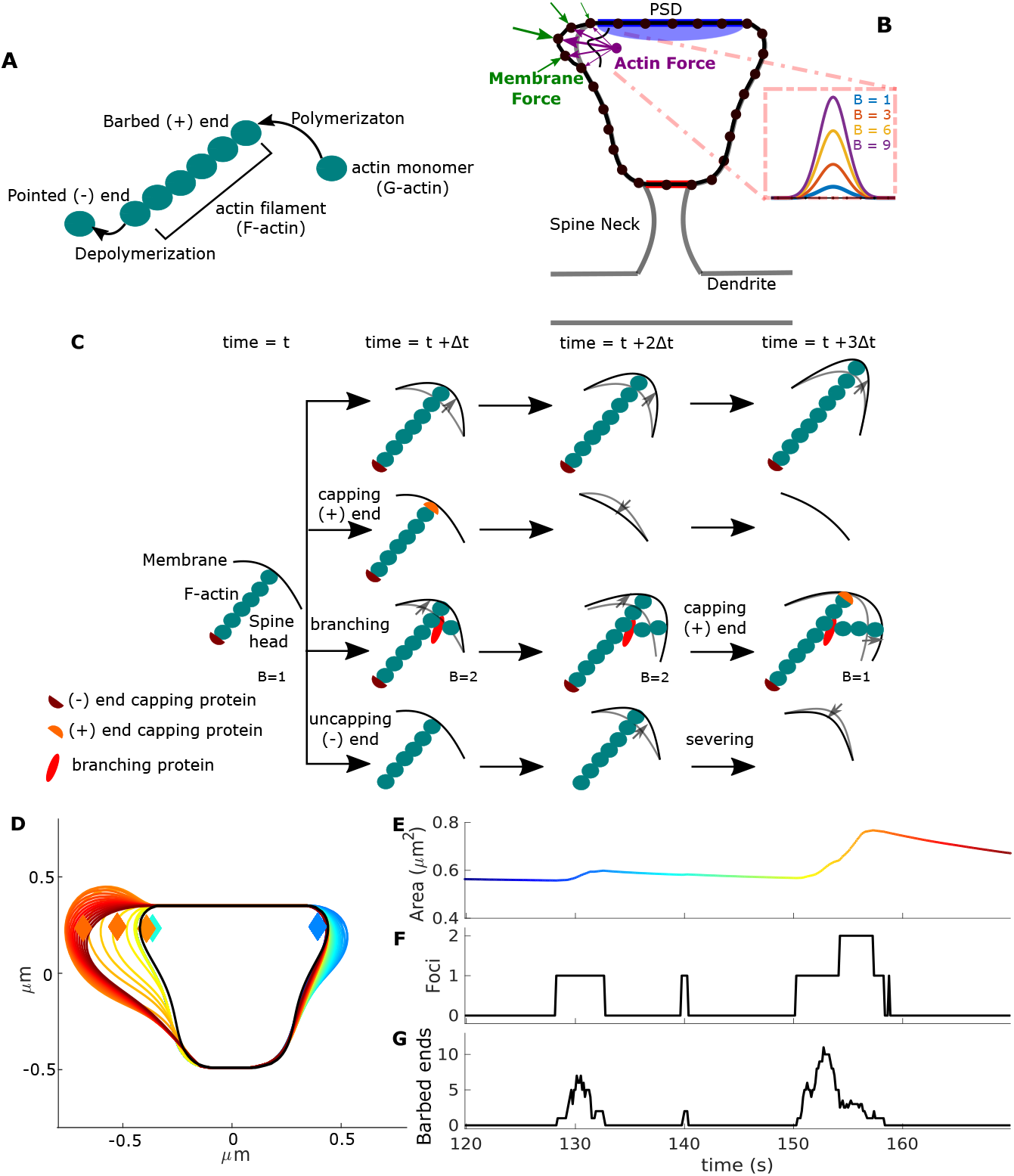
Theoretical model: Actin-Membrane interactions. The mechanisms that displace and change PSD and spine neck are not shown in this figure. **A** Actin treadmilling process. **B** Filament events for 4 time-steps, each row shows a different scenario. At each time-step, all filaments with uncapped (+) end undergo monomer addition. Also, uncapped (+) ends can branch with certain probability, hence *B ⟶ B* + 1, where *B* is the number of uncapped (+) ends. Additionally, there is a probability that a capping protein attaches to an uncapped (+) end then preventing actin polymerization, hence *B ⟶ B* − 1. When there are no filaments with uncapped (+) ends at a focus, it vanishes. In addition, a capped (-) end can become uncapped and an uncapped (-) end can be severed (thus, *B ⟶ B* − 1). **C** Force interactions in a dendritic spine head. **D** Spine shapes arising from model simulation, color-code according to time as in (E). Black line represents the “resting” shape to which the spine shrinks in the absence of actin force. Diamonds represent actin nucleation locations, initiated at different times (color-coded). **E** Area evolution of the spine, color-coded from initial (in dark blue) to final time. **F** Number of polymerization foci during this period. **G** Number of uncapped (+) ends at the polymerization foci over time. See Methods for further details.

These different mechanisms lead to spine shape fluctuations that mainly arise from the continuous nucleation as well as from the disappearance of actin polymerization foci at different locations. Therein, the actin polymerization process yields changes of the number of barbed ends at each time-step, which alters the force generated by actin; and thus, the counter-force generated by the membrane (see Figs. 4D-G).

### Comparison between model simulation and experimental data

Figure 5A shows the resulting spines shapes. To determine if the model exhibits similar shape fluctuations than the experimental data, we divided the simulation into 5-minute time windows with the spine shapes sampled every 10 seconds and calculated the area and shape descriptors (Fig. 5D-G). Shape descriptors are different for each 5-minute time window of the simulation (Fig. 5C). Note that this variability is the result of sustained fluctuations over time (Fig. 5D-G), rather than of a large single change, hence the variability over time is maintained (Fig. 5H). Moreover, the spine area and its changes are significantly correlated to those in *S* (Fig. 5I-N).

**Figure 5:**
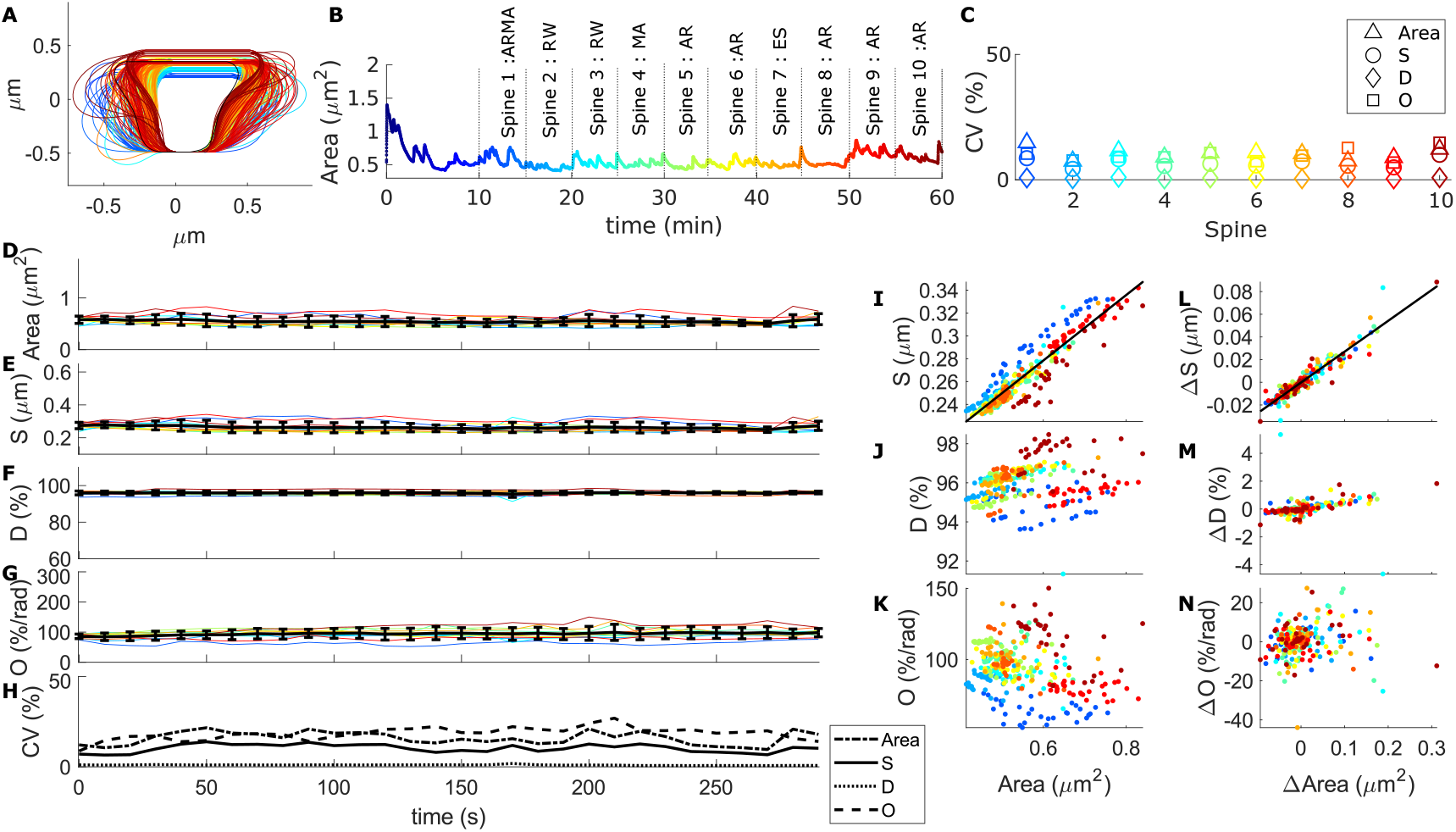
Simulations. **A** Spine shapes resulting from the model simulation every 10 seconds, color-coded from 0 minute (dark blue) to 60 min (dark red). **B** Spine area evolution over time corresponding to the simulation in A, dotted lines indicate the 5 minute windows, the spine number and the best fit of an ARIMA model (abbreviations as in Table S1). **C** Coefficient of variability of the shape descriptors for each time window. **D-G** Shape descriptors over time for the 5 minute windows (spines in C) of the simulation. As in the experimental data, the values were sampled every ten seconds. **H** Coefficient of variability of shape descriptor evolution over time. **I-K** Spine area against *S*, *D* and *O*. Each dot represent the value for a certain time point and color-code corresponds to the respective 5 minute window. Black lines correspond to a linear least-squares fit with a significant Pearson’s correlation coefficient of 0.93 in I (p-value *<* 0.01 using a Student’s *t* distribution). **L-N** Difference in spine area against Δ*S*, Δ*D* and Δ*O*, respectively. L has a significant Pearson’s correlation coefficient of 0.95.

To more directly compare model results to experimental data, we ran ten 60-minute simulations. Figure 6 shows that the model reproduces the same shape characteristics as the experimental data. The difference between the shape descriptors’ CVs of the model and the experimental data can be due to inaccuracies in the extracted ROIs. Thus, instead of calculating the descriptors from the original simulation spine images, we calculated them from modified model images that better resemble confocal microscopy ones. Such images, obtained by randomly allocating fluorophores inside the modeled dendrites and blurring them, were analyzed like experimental data on Fiji. In the Supporting Information we provide more controls of this kind showing that the variability of the shape descriptors is increased and, this way, matches the higher variability of the experimental data. Therefore, the difference in CVs between experimental data and model is very likely due to inaccuracies in obtaining a ROI from experimental data. Moreover, the time series analysis of the 5-minute time windows show that returning to a constant mean is the most common trend (40% of the time windows).

**Figure 6:**
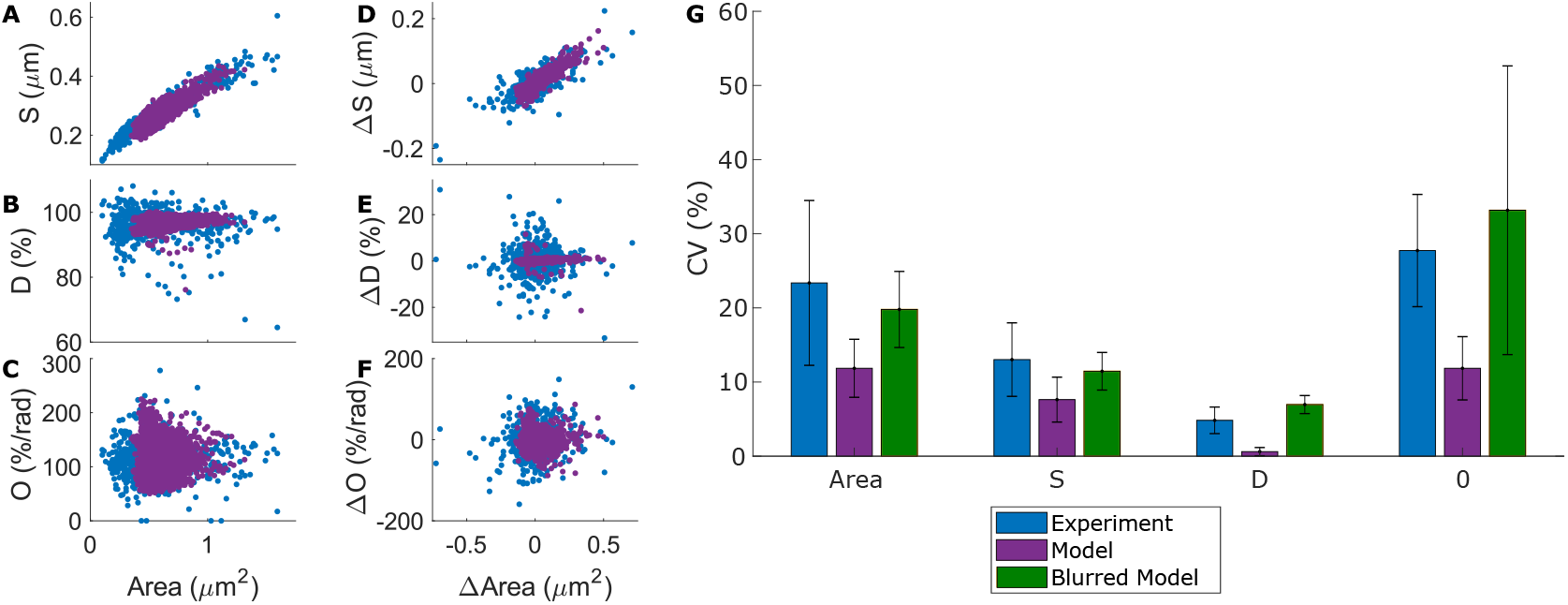
Comparison between experimental data and model. **A-C** Spine area against shape descriptors *S*, *D* and *O*. Each dot represents a data point at a sampled time frame. There are 16 spines sampled every 10 seconds for 5 minutes corresponding to experimental data from Figure 2 (in blue) and frames for the model (purple) correspond to 10 simulations in which shapes are sampled every 10 seconds from minute 10 to minute 60. **D-F** Changes in shape descriptors from the data in (A-C) against changes in area. **G** Median ± standard deviation of the coefficient of variability for shape descriptors calculated for each spine, in experimental data or in the model. Green bar corresponds to the CV’s of the model sampled shapes, but blurred.

### Presence of 1*/f* noise

Note that the area evolution within every 5-minute time window in the model shows different tendencies, even for consecutive windows; however, returning to a constant mean is the most common trend (AR in Fig. 5B), as in the experimental data. Thus, for this type of behavior we investigated, using our model, whether a general trend would emerge for longer durations, which exceed those currently accessible by experiments. We first observed that the area evolution corresponding to the sampled points from Figure 5B sometimes fluctuates around low values and other times around high values. However, there is no sustained increase or decrease of the mean and standard deviation of the area, or any cyclical trend, i.e., the time series is stationary (according to an augmented Dickey-Fuller test with p-value *<* 0.01). Further analysis shows that the autocorrelation function of these samples points follows a slow asymptotic decay (Fig. 7A), which is a feature of 1*/f* noise. The main characteristic of this type of noise is that it arises from a stochastic process with spectral density *SD*(*f*) that follows a power law, i.e., *SD*(*f*) = 1*/f ^α^*, with frequency *f* and power law exponent 0.5 ≲ *α* ≲ 1.5.

**Figure 7:**
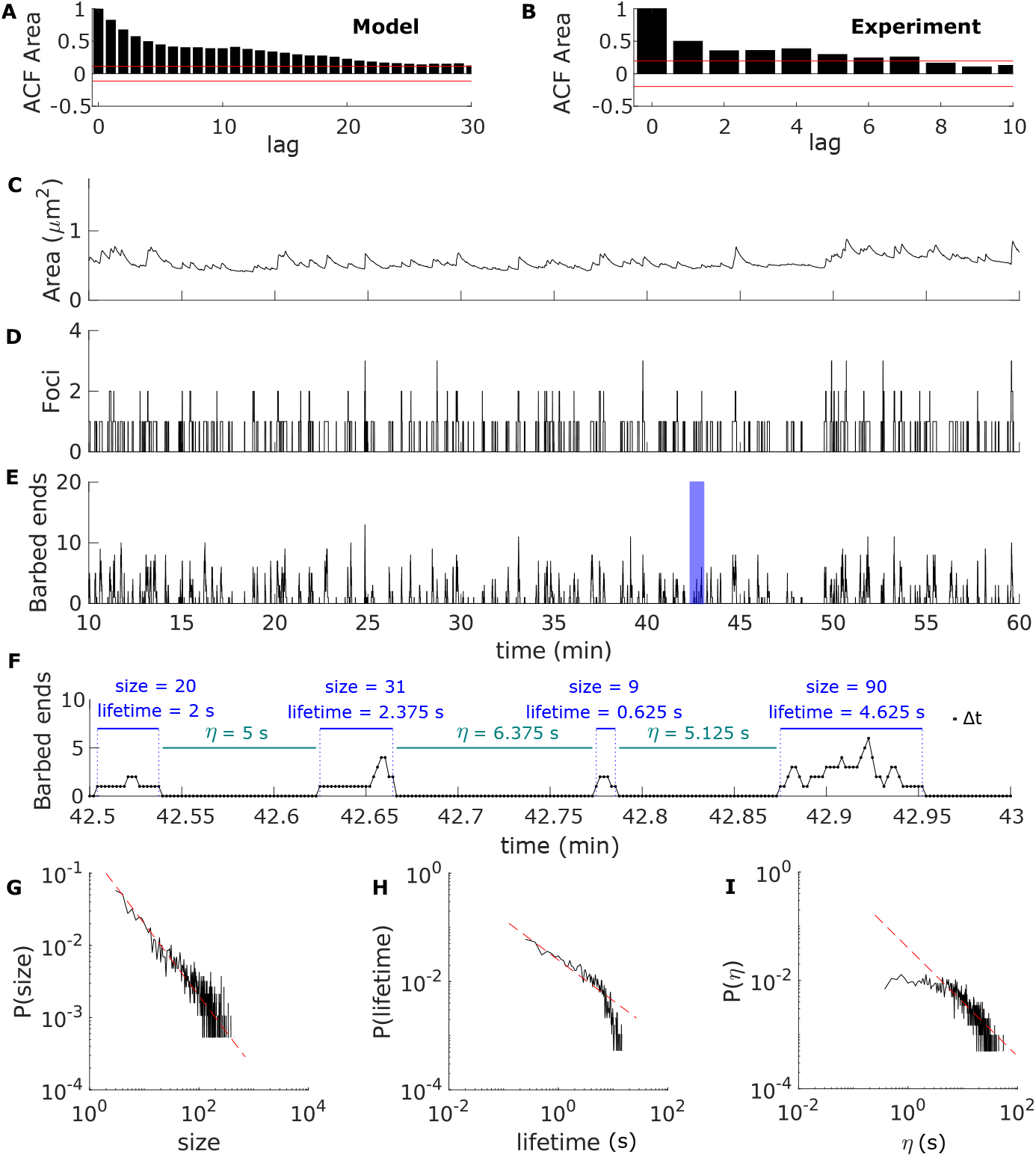
Self-organized criticality. **A** First lags (10%) of the autocorrelation function of the sampled area from Figure 5B. Dashed red lines correspond to the 95% confidence bands. **B** Same as (A) but calculated from the experimental data in Figure 3A. Spine area **C**, number of actin polymerization foci **D**, and total number of barbed ends in the spine **E** corresponding to the simulation in Figure 5. **F** Zoom of the blue-shaded interval in (E). Each dot indicates the number of barbed ends at each time-step. The size and duration of the avalanches of actin polymerization, as well as the interval between avalanches (*η*), is displayed. Log-log plot of the probability of avalanche size **F**, lifetime **G**, and duration of the interval between avalanches **I** corresponding to the data from 10 simulations of the model. Red dotted lines correspond to straight lines with slope of −1 (D and F) and −1.36 (E), plotted for visualization purposes.

To test the presence of 1*/f* noise in the time series corresponding to the spine area sampled every 10 seconds from minute 10 to minute 60 (i.e., assembling all 5-minute time windows of the simulation), we used the method developed by Wagenmakers et al. (2004). This method is based on the idea that 1*/f* noise (related to long term correlations) appears between white noise and a random walk (see Methods for details).

When applying this method to the sampled area trace from Figure 5B, we verified that the time series indeed behaves like 1*/f* noise. In addition, we tested each of the ten simulations of the model from Figure 6 (in purple) for the presence of 1*/f* noise, and we found that it is present in 80%. Furthermore, we found that the time series of the spine area from the longer experimental recordings (Fig. 3A) is also stationary (according to an augmented Dickey-Fuller test with p-value *<* 0.01) and the autocorrelation function decays asymptotically (Fig. 7B). Also for this data set the method by Wagenmakers et al. (2004) confirms that the spine area evolution is dominated by 1*/f* noise.

### Self-organized criticality explains *1/f* noise in spine size fluctuations

To explain the presence of 1*/f* noise in the proposed model, we explored which mechanism could be generating it. Perhaps one of the most common theories, proposed by Bak et al. (1987), is that 1*/f* noise emerges when a system is in a self-organized critical state where small fluctuations cause events of all sizes with a probability density function that is described by a power law. The durations of these events are related to their size; and hence, described by a power law function that correlates to 1*/f* noise. Therefore, this theory connects the temporal 1*/f* noise with the evolution of a spatial structure with scale-invariant, self-similar properties (Bak et al., 1988).

In our model, the changes in spine size are due to the force exerted by polymerization of actin barbed ends at distinct foci. Hence, the fluctuations in spine area correlate to the number of barbed ends (significant Pearson’s correlation coefficient of 0.6751 with p-value *<* 0.01 using the data from Fig. 7C and E). Polymerization foci nucleate with a given probability at random locations, preferably located close to the PSD, and have finite lifetime, due to the extinction of barbed ends. At a given focus, the number of barbed ends fluctuates at each time-step due to capping, branching and severing events (see Fig. 4). Importantly, the branching probability depends on the membrane force and number of barbed ends in the foci. Therefore, it acts as a feedback mechanism that decreases the probability of branching if there are many barbed ends in the focus or if the membrane force increases. The latter is the result of the local curvature increase of the spine membrane due to the protrusion formed by the actin force. Figure 7E shows that the total number of barbed ends in our simulations behave like avalanches, in the sense that they have finite duration and different lengths and sizes. Hereinafter, we define the size of an avalanche in our system by the number of polymerization events. Since the barbed ends polymerize at each time-step, size is equivalent to the sum of barbed ends at each time-step during the avalanche (see Fig. 7F).

To examine whether our model has spatio-temporal power law characteristics, we collected all avalanches for ten 60-minute simulations (corresponding to Fig. 6) and calculated the probability of their size and duration. Figure 7G-H show the log-log plot of such probabilities, which are for long stretch straight and have negative slope. Hence, they behave like power law functions. The deviation from the straight line in Figure 7H is due to the finite lifetime of the polymerization foci (finite-size effect in Bak et al. (1988)). Interestingly, as in Bak et al. (1987), the mean of the 1*/f* noise exponent resulting from the ARFIMA fittings *α ≈* 0.72 relates to the exponent of the power law of the event lifetime probability distribution 1.36 *±* 0.008 *≈* 2 *− α*, calculated as in (Newman, 2005). Moreover, the intervals between avalanches follow a power law (see Fig.7I). These features also arise in simulations with different sets of parameters (see Supporting Information). Moreover, various other tests confirm that our simulations show criticality (see Supporting Information). Therefore, we concluded that actin polymerization in the modeled dendritic spines self organizes to a critical state.

## Discussion

In the current study we investigated the function of spontaneous shape fluctuations in dendritic spines in time-lapse experimental data using novel quantification methods. To examine the behavior of spines over longer timescales, we performed simulations of a biophysical model that mimics spine shape fluctuations based on actin dynamics. Shape fluctuations in the model follow 1*/f* noise and the model predicts that this might be due to a process of self-organized criticality of the actin dynamics in a spine.

### New Analysis Methods to Characterize Spine Shape and Size

When studying fluctuations of a signal over time, the distinction between relevant information and noise is crucial. Here, we used time series analysis to detect whether there are certain characteristic tendencies in the dendritic spine size changes over time. Specifically, we used ARIMA models to analyse the experimental data. We found that the behavior of dendritic spine sizes in 5-minute recordings shows diverse trends, albeit returning to a constant mean, captured by an ARIMA(1,0,0) model, was the most common behavior. This novel approach appears, thus, promising for obtaining additional quantitative information from experiments by creating forecasts of the time series behavior over longer temporal periods (Hyndman & Athanasopoulos, 2018; Box et al., 2015).

We observed that the shapes of dendritic spines are highly asymmetric, and hence, often employed, common shape factors; such as the circularity index or aspect ratio measures; fail to describe this well. For example, the circularity index describes how much the shape deviates from a circle, but it cannot capture shape elongations. On the other hand, the aspect ratio characterizes shape elongations but fails to distinguish *different* elongated shapes (Wojnar et al., 2000). Therefore, in order to better describe the evolution of spine shapes, we proposed new set of shape descriptors. Such descriptors are based on concepts of circular statistics (Batschelet, 1981) that have been implemented, for example, to describe the asymmetry of visual cortical response curves to moving, oriented stimuli (Wörgötter & Eysel, 1987; Li et al., 1994).

We adapted these descriptors to characterize the directional and orientational selectivity of the spine shape and found that they fluctuate randomly over time in most of the cases. Moreover, using these new shape descriptors we showed that, in line with the previous findings of Fischer et al. (1998), shape changes are uncorrelated to size changes in dendritic spines. Importantly, this holds also for a second, different data set verifying the general character of this property.

### Modelling Spine Size and Shape Dynamics

In general, all experimental data sets are still rather short and the long-term temporal evolution of a spine cannot yet be very well measured. To overcome this limitation, we implemented a theoretical model, based on (Bonilla-Quintana et al., 2020), and showed that it captures the experimental data well. The model contains two central features: 1) actin treadmilling dynamics that arises from few, short-lived actin foci and 2) much slower random processes that allow spine neck and PSD fluctuations. Such fluctuation have been observed in the here-analyzed data and also in other studies (Tønnesen et al., 2014; Minerbi et al., 2009). In this way, the model is compatible with physiologically relevant findings at individual spines, and also in the neuronal network, that show the importance of PSD changes during the lifetime of a spine (see Ziv & Brenner (2018) for a review).

### Spine Size Over Long Periods Exhibit *1/f* Noise

We used our model to explore the behavior of spine size fluctuations for long durations and found the presence of 1*/f* noise. Moreover, we showed that 1*/f* noise is also present in longer experimental recordings. However, more experiments are needed to consolidate this finding, which is currently supported by only one data set. Interestingly, this type of noise has been previously identified in the brain at quite different temporal scales. For all of the following phenomena there is evidence for the presence of 1*/f* noise: Spike discharge intervals in tonically activated neurons (Musha & Yamamoto, 1997), action potentials of a squid giant axon (Musha et al., 1981), voltage time course of nerve cell membranes (Verveen & Derksen, 1968; Lundström & McQueen, 1974), the EEG alpha rhythm (Musha & Yamamoto, 1997). Futhermore, also in a human cognitive temporal estimation task, 1*/f* noise has been found (Wagenmakers et al., 2004). Therefore, 1*/f* noise seems an ubiquitous feature of the nervous system.

### Actin in the Spine Self-organizes into a Critical State - Suggested Experiments

As a consequence, we explored which mechanism could generate 1*/f* noise in spines. Although the concept of self-organized criticality and its role in the brain has been controversial since its origin (Watkins et al., 2016; Beggs & Timme, 2012), we chose this concept with the aim to link spatial and temporal phenomena in our model, as in the original interpretation of Bak et al. (1987). Because blockage of actin polymerization in spines inhibits spine shape changes (Fischer et al., 1998; Dunaevsky et al., 1999), we proposed actin polymerization as the molecular mechanism that causes 1*/f* noise. Hence, we studied the size and length of polymerization events in our model and found that they exhibit robust spatiotemporal power law correlations. Since this feature does not depend on the fine tuning of model parameters (see Supporting Information), we concluded that in the proposed model actin dynamics self-organize into a critical state.

The model predicts that there are polymerization *avalanches*, which occur in bursts originating from different foci. This suggests future experiments, proposed below, to confirm or disprove this prediction. Currently there is, however, some indirect evidence that supports this.

For example, Honkura et al. (2008) detected two distinctive pools of actin, a static and a dynamic one within the spine and Frost et al. (2010) observed that fast polymerization, corresponding to the dynamic pool, occurs at discrete foci. Thus, there is always a certain equilibrium existing between free actin monomers and actin filaments (F-actin and G-actin) in the spine. For example, Okamoto et al. (2004) observed that this equilibrium *globally* changes during spontaneous width changes of dendritic spines.

The existence of discrete polymerization foci (Frost et al., 2010) shows that the equilibrium is also often *locally* disturbed by the gathering of monomers that bind and start a new filament. However, the amount of locally accessible free actin is finite, and hence, the lifetime of any focus and its filaments is limited (also due to capping processes, etc.). These experimentally confirmed processes can, thus, create the balance between new actin filaments outgrowth and their collapse required to bring such a system into a self-organized critical state. We speculate that this creates a system, which is highly susceptible to change and can react efficiently for example when a spine will grow quickly following, for example, the induction of long term potentiation (Bosch et al., 2014; Chazeau & Giannone, 2016).

While these observations are phenomenological, the model suggests a possible biophysical mechanism that could underly this. In general, a critical state is maintained by a constant flow of energy and material that is reached via feedback mechanisms (Mora & Bialek, 2011). Self-regulation in the model occurs through a feedback mechanism in the branching rate of the barbed ends, which depends on the local shape configuration and the number of barbed ends at each actin polymerization focus. Thus, in our model, global criticality emerges from the local properties of the different foci. This leads indeed to a balance between static filaments and free actin monomers.

To validate this hypothesis, new experiments measuring the size and duration/lifetime of the actin polymerization foci for long timescales are needed. Such experiments are very challenging since the actin structure within a spine is dense, leading to signal saturation and difficulties in the visualization of single filaments even with the most advanced nanoscopy techniques. Furthermore, imaging at high spatial and temporal resolution for long timescales is impaired by the bleaching of the reporter molecules. Lastly, labeling strategies relying on the direct labeling of endogenous actin (e.g. SiR-Actin, see Lukinavičius et al. (2014)) are based on Jasplakinolide and hence might interfere, even if only minimally, with the fine regulation of actin dynamics in the spines.

Hence, an experiment to validate our hypothesis should aim at labeling only a sparse subset of actin, thereby enabling the visualization of small changes in concentration. Besides fluorescence intensity, another possible readout is FRET, that could allow discriminating between F- and G-actin (Okamoto et al., 2004). The sparse labeling could be achieved by short pulse-chase experiments in cells expressing actin in fusion with a self-labeling tag (e.g SNAP-Tag or HaloTag, Hoelzel & Zhang (2020)) or by injecting a relatively low amount of labeled actin monomers. The optimal image acquisition rate for longer experiments can be estimated after studying the focus lifetimes over short periods of time but at high frequency.

An estimate of the number of polymerization events at a given time can be achieved by super resolution multi-color imaging of all the actin filaments, which can be labeled by phalloidin, of capping proteins that inhibit filament (+) end growth, such as CapZ, and of proteins present where the filaments start at branching points, such as Arp2/3. The total number of polymerizing events can be approximated by counting the filaments close to the membrane without CapZ. However, this is only possible in fixed samples, and thus, the fluctuations of barbed ends over time cannot be captured.

Addressing the role of potential actin avalanches experimentally may, thus, shed a light on the question whether spontaneous spine size- and shape fluctuation is an epiphenomenon or a reflection of a functionally relevant state - as suggested by this study.

## Acknowledgements

We thank the German Science Foundation through the Collaborative Research Center SFB-1286, Projects C3 (MB-Q,FW) and A7 (ED), for funding this research. We are grateful to Dr. Marina Mikhaylova for kindly providing her experimental data. We thank her and the members of the SFB-1286, especially Silvio Rizzoli, for fruitful discussions.

## Author Contributions

MB-Q, FW, CT and MF contributed to the study concept. MB-Q performed model simulations and formal analysis, and wrote the original draft. ED performed experiments. MB-Q, FW, CT and MF analyzed the results. All the authors edited the paper.

## Methods

### Key Resources Table

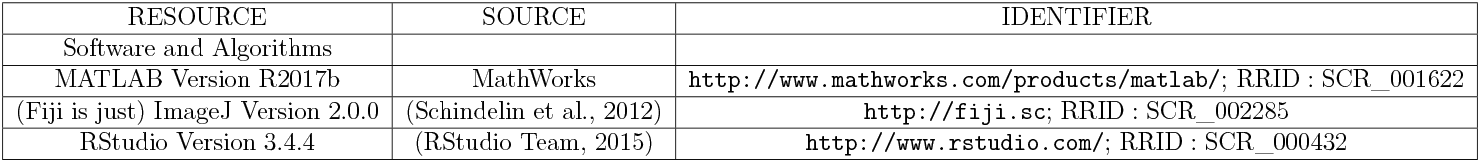

| RESOURCE | SOURCE | IDENTIFIER |
| --- | --- | --- |
| Software and Algorithms |  |  |
| MATLAB Version R2017b | MathWorks | http://www.mathworks.com/products/matlab/; RRID : SCR_001622 |
| (Fiji is just) ImageJ Version 2.0.0 | (Schindelin et al., 2012) | http://fiji.sc; RRID : SCR_002285 |
| RStudio Version 3.4.4 | (RStudio Team, 2015) | http://www.rstudio.com/; RRID : SCR_000432 |

### Experimental Data Analysis

We analyze experimental data provided by Dr. Mikhaylova that corresponds to Figures 6 and S7 in (Mikhaylova et al., 2018) and consists of time-lapse confocal microscopy images taken every 10 seconds for 5 minutes from wild type mouse hippocampal primary cultures at 12 days in vitro (DIV12) co-transfected with mRuby2 (volume marker) and GFP-actin.

We also analysed a different data set from the lab of E.D., which consist of 100 frames taken every 9.6 seconds. For this, cultured hippocampal neurons at 19 DIV were stained with 2 *μ*g*/*mL DiO (Thermo Fisher) for 20 min in ACSF, washed for 10 minutes and imaged in confocal mode in ACSF buffer at room temperature.

We process these images using Fiji (Schindelin et al., 2012). First, we trace a square region of interest (ROI) around a spine and duplicate the entire stack of images in the GFP-actin channel. We apply a Kuwahara Filter with a kernel size of 3, which preserves edges; and, thus, the small protrusions of the spines. The resulting image is thresholded (using the default method) and then a mask is created in which a watersheed segmentation is applied to separate the spine head from the neck. Finally, a ROI is created around the spine using the Analyze Particles function and the neck position is manually traced (see Fig. 1A). Subsequently, we load the ROIs into Matlab using the function ReadImageJROI.m from (Muir & Kampa, 2015).

### Obtaining *R*(*θ*)

Following Li et al. (1994), to obtain *R*(*θ*) in Eq. (1) (orange dotted line in Fig. 2B), we perform an optimal periodic regression on dROI(*θ*) (brown line in Fig. 2B). First, an “entire fitting” of dROI(*θ*) is computed by

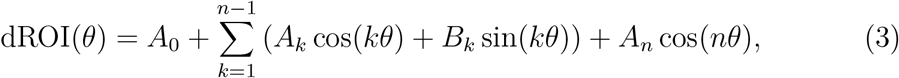

where the regression coefficients (*A_k_*, *B_k_*) are given by

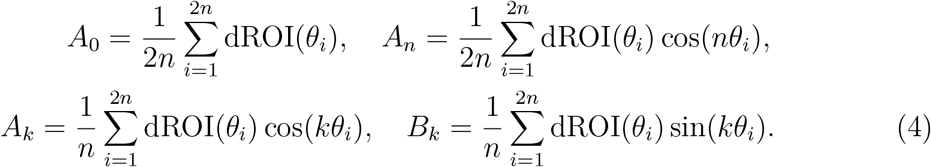

Here, *i* = 1, 2*, …,* 2*n* is the number of sampling points corresponding to the angle *θ_i_* = (*i −* 1)*π/n*, with *n* = 12. As in Li et al. (1994), we assume that an entire fitting to the data is improper due to the errors that can arise from the experiment or the data processing. Therefore, all the terms with a standard regression coefficient smaller than 0.1 are discarded (one by one) if they do not significantly alter dROI(*θ*) (assessed using a F-test with a level of significance of 0.1). The resulting function *R*(*θ*) is periodic and, hence, can be divided in three parts:

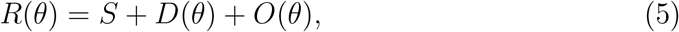

where

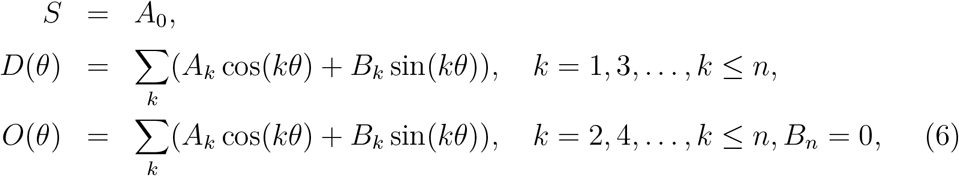

and the discarded terms are set to zero (Fig. 2C).

### Theoretical Model

We are basing our model here on the 2D model with fixed PSD and neck described in (Bonilla-Quintana et al., 2020). Note that in the older study a detailed verification against simulations in 3D (full spine volume mesh-model) is provided to show that the 2D reduced model captures the actin-membrane interaction accurately. We first describe the main characteristics of the basic model (for all details see Bonilla-Quintana et al. (2020)) and then the extensions made for the current study.

The spine membrane is approximated by a mesh Γ of *n* vertices at position **x**^*k*^ = (*x^k^, y^k^*) *∈* ℝ^2^, *k* = 1*, …, n*. At every time-step:

- A polymerization focus *i* nucleates at a random location 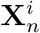 (close to the PSD) with probability *γ_f_* Δ_*t*_.
- Each barbed end of F-actin polymerizes G-actin. Additionally,

- Uncapped barbed ends branch creating a new filament with a probability Δ_*t*_*γ*_*branch*_(*t*).
- Uncapped barbed ends cap with a probability Δ_*t*_*γ*_*cap*_. Consequently, the barbed end stops generating force and, hence, the corresponding filaments are eliminated from the simulation.
- Capped minus ends uncap with a probability Δ_*t*_*γ*_*uncap*_.
- Uncapped minus ends are eliminated with a probability Δ_*t*_*γ*_*sever*_.
- Mesh vertices displace according to

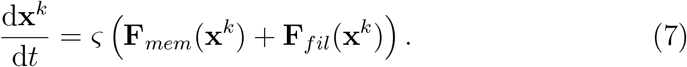

In Eq. (7) *ζ* is a proportionality constant, **F**_*fil*_ and **F**_*mem*_ are the actin and membrane forces, respectively. Here

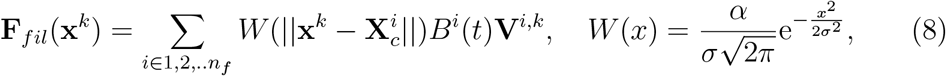

where α denotes the amplitude and *σ* the standard deviation of the Guassian kernel *W*, *n_f_* is the number of active actin foci with *B* barbed ends, and 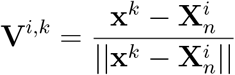 the normalized direction vector of the force from focus *i*, located at 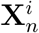.

The membrane force is given by

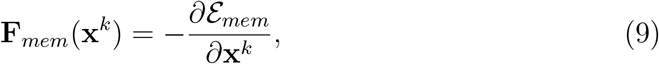

where

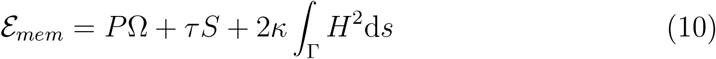

is the Helfrich free energy. Here *P* is the difference between internal and external pressure, *τ* the line tension, and *κ* the bending modulus. Ω denotes the area enclosed by the membrane, *S* the boundary length, and *H* the mean curvature.

All the rates are constant, except the branching rate

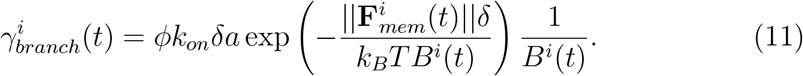

that changes at every time-step and depends on the number of barbed ends at each focus *i*, *B^i^*, and the membrane force. Here, *T* is the absolute temperature, *k_B_* the Boltzmann’s constant, *δ* the length of an actin monomer, *k_on_* the barbed end monomer assembly rate, *ϕ* the branching rate amplitude and *a* the concentration profilin-ATP-actin available for polymerisation. All parameters are the same as in (Bonilla-Quintana et al., 2020), except for *σ* = 0.2 (see Table S3).

This basic model has been extended here to incorporate the fact that the PSD and the neck can change (on a slow timescale), too. Thus, the PSD lateral movements and size changes used in the current study are implemented by introducing four random processes:

- The PSD right boundary elongates (moves to the right) with probability Δ_*t*_*γ*_*r*+_.
- The PSD right boundary shrinks (moves to the left) with probability Δ_*t*_*γ*_*r−*_.
- The PSD left boundary shrinks (moves to the right) with probability Δ_*t*_*γ*_*l−*_.
- The PSD left boundary elongates (moves to the left) with probability Δ_*t*_*γ*_*l*+_.

For this, we draw a number from a uniform distribution for each process. If the number is less than the designated probability, then the PSD elongates and/or shrinks, respectively. Such events are implemented by adjusting the vertices in the spine mesh corresponding to the PSD. When the PSD right (left) boundary shrinks, the first (last) vertex corresponding to the PSD is released, in the sense that no longer forms part of the PSD and can move according to the forces generated by actin and the membrane. When the PSD right (left) boundary elongates, the PSD incorporates the neighboring vertex to the right (left) from the spine. Consequently, the vertex is fixed with the *y*-coordinate set to *h_PSD_*. Note that these events are independent, and thus, can result on a lateral displacement of the PSD with respect to the neck.

Likewise, the PSD moves vertically upward by setting *h_PSD_* = *h_PSD_* + Δ*d_mov_* with probability Δ_*t*_*γ*_*up*_, and moves vertically downward decreasing *h_PSD_* by Δ*d_mov_* with probability Δ_*t*_*γ*_*down*_. If the PSD moves vertically, all the corresponding vertices in the mesh change their *y*-coordinate to the new value of *h_PSD_*.

Since the neck center is the reference point to calculate the here-proposed shape descriptors, we prefer to avoid its displacement by only allowing symmetrical neck width changes. For this, we elongate the neck with probability Δ_*t*_*γ*_*n*+_ by recruiting the right and left neighboring spine vertices and fix them to *h_neck_*, or shrink the neck with probability Δ_*t*_*γ*_*n−*_ by releasing the initial and final vertices corresponding to the neck.

Note that all these events are independent. We select *γ*_♦_ = 1*/*75 s^*−*1^, ♦ *∈ {r*+*, r−, l*+*, l−, n*+*, n−, up, down}*, which renders a good qualitative fit to the data. For *d_mov_*, we use the same value as the minimum distance allowed between mesh vertices *d_min_* = 0.018 *μ*m. In the simulation, the changes to the PSD and neck are computed after the actin dynamics.

### Quantification and Statistical Analysis

The tendency analysis on the spine area and shape descriptors is performed in RStudio using the `auto.arima()` function with `max.p=1`, `max.q=1`, `max.d=1`, `start.p=0`, `start.q=0`, to ensure the selection of simple models, and assessed by the Akaike information criterion (AIC). We test if the residuals of the selected model are significantly stationary according to the augmented Dickey-Fuller test (`adf.test()` with 0 lag and p-value *<* 0.01). Moreover, the residuals’ autocorrelation (ACF) and partial autocorrelation (PACF) function are plotted. If the residuals of the selected model are non-stationary and/or have pronounced peaks in the ACF or/and PACF, we investigate which of the possible 8 ARIMA models best fit the data using the `arima()` function and AIC.

To test the presence of 1*/f* noise in the time series corresponding to the spine area, we use the method developed by Wagenmakers et al. (2004). This method is based on the idea that 1*/f* noise (related to long term correlations) appears between white noise (ARIMA(0,0,0)) and a random walk (ARIMA(0,1,0)). Note that white noise, denoted by *X_t_* = *E_t_*, where *X_t_* is the value of an observation at time *t* and *E_t_* denotes a random number drawn from a normal distribution, is obtained taking the difference between successive observations in a random walk, given by *X_t_* = *X_t−_*_1_ + *E_t_*. This is capture by the parameter *d*, which denotes the number of difference needed for stationarity, in their corresponding ARIMA(*p, d, q*) model. Moreover, the spectral density of a random walk has a power law exponent of *α* = 2, and the white noise has *α* = 0, thus *d* = *α/*2. This implies that, to model 1*/f* noise, *d >* 0 must include fractional numbers, as in ARFIMA (autoregressive fractionally integrated average) models.

Following Wagenmakers et al. (2004), we test whether the spine area time series presents long-range or short-range dependencies, i.e., whether the time series fits better an ARFIMA(1,*d*,1) or an ARIMA(1,0,1) model. We apply the `fracdiff()` function in RStudio with `nar=1`, `nma=1`, to get the coefficients of the ARFIMA(1, *d*,1) model, and check if they are significantly greater than zero using `summary()` with `signif.stars = TRUE`. A time series exhibits 1*/f* fluctuations if *d* is significantly different from zero and if the AIC of the ARFIMA(1,*d*,1) model is greater than that of an ARIMA(1,0,1) model. For calculating the AIC of the ARIMA(0,1,0) model, we use the arima function. The residuals of both models are tested for stationarity as above.

## Data and Code Availability

### Data availability

Data used to generate the findings of this study will be freely available upon request. The data corresponding to Figure 1 has been previously published in (Mikhaylova et al., 2018) and it is available from there.

### Code availability

Custom computer code used to generate the findings of this study will be made available upon request.

## Supporting Information

### ARIMA fittings

**Table S1:** ARIMA FITTINGS

| Model | Area | S | D | O |
| --- | --- | --- | --- | --- |
| Spine 1 | WN | WN | WN | WN |
| Spine 2 | DA | ES | AR | ES |
| Spine 3 | DA | DA | WN | WN |
| Spine 4 | MA | MA | AR | AR |
| Spine 5 | AR | AR | MA | WN |
| Spine 6 | RW | RW | WN | ES |
| Spine 7 | ES | RW | WN | WN |
| Spine 8 | ES | ES | MA | AR |
| Spine 9 | ES | WN | WN | WN |
| Spine 10 | AR | WN | MA | WN |
| Spine 11 | MA | ES | WN | ES |
| Spine 12 | AR | AR | ES | ES |
| Spine 13 | WN | WN | WN | MA |
| Spine 14 | DA | DA | WN | MA |
| Spine 15 | AR | RW | ES | WN |
| Spine 16 | AR | AR | RW | WN |

### Shape Descriptors

In the following shape descriptors are interpreted. To achieve this, five different kinds of spine heads are depicted (left column of Fig. S1) and their shape descriptors are plotted (right column of Fig. S1).

#### *S* corresponds to spine head size

Note that *S* corresponds to the first term of *R*(*θ*) in Eq. (1) (see Fig. S1, center column, dashed line). Since *R*(*θ*) is periodic, *S* represents the average value of the function over all the domain (*θ ∈* [0, 2*π*)). Hence, *S* gives an average of the distance from the spine neck center to the membrane of the spine head at the sampling points. In this way, *S* can be used as an indicative of the spine head area. Moreover, there is a correspondence between area and *S* values in the sample spines: bigger spines have larger *S* value, and vice versa.

#### *D* indicates direction selectivity

Note that *D*(*θ*) in Eq. (6) is an odd function (see Fig. S1, center column, dotted-dashed line). For a “normal” spine shape (Fig. S1 A), the maximum distance from the spine neck is assumed to be aligned with the neck center. Thus, the maximal value of *D*(*θ*) is *≈ θ* = *π/*2. Moreover, the shape of *D*(*θ*) is similar to a sinusoidal function with period 2*π* and amplitude *≈ πS/*2. Therefore, when *D*(*θ*) is integrated and averaged (see Eq. (6)) it gives *D ≈* 100% of *S*.

Importantly, when the spine head shows a clear preference for a direction from the spine neck, *D*(*θ*) has a clear peak that resembles a sinusoid and, thus, *D ≈* 100% of *S* (see Figs. S1 A-C). If all the sampling points are equidistant to the spine neck (Fig. S1 D) then the amplitude of *D*(*θ*) is close to *S*. Hence, *D <* 100% of *S*. Likewise, *D <* 100% of *S* if the spine head has various bumps, which have similar distance from the neck (Fig. S1 E). In this case, *D*(*θ*) peaks at the bump locations; however, these peaks are around *S* and the integration results in a smaller percentage of *S*. Hence, *D* is an indicator for spine head preference of direction from the spine neck. If the spine is centered or leans towards a side, then *D ≈* 100%. If the spine head membrane is equidistant to the neck or if the spine head has several bumps, then *D <* 100%.

#### *O* indicates orientation selectivity

*O*(*θ*) is an even function (see Fig. S1, center column, solid line). Accordingly, only half of the period is considered when calculating *O* (see 6). Note that *O* is the averaged integral over the derivative of *O*(*θ*) (see 2). Therefore, it represents the changes in the slope of *O*(*θ*) between sampling points. For example, for a “normal” centred spine head shape (Fig. S1 A), *O*(*θ*) is close to a cosine function with period 2*π* and amplitude 2*S*. Thus, the changes of its slope between sample points have to be of the same magnitude and *O ∼* 4*S/π*.

When the spine head is more elongated with some orientation (Figs. S1 B-C), then the slope of *O*(*θ*) is steeper and *O >* 400*/π*. For sharper elongations, *O* is larger (Fig. S1 B). Likewise, when there are several sharp protrusions (Fig. S1 E) the value of *O* is larger. Finally, if the spine head does not have any elongation (Fig. S1 D), then *O*(*θ*) is almost constant and consequently, *O* is small. Therefore, *O* denotes spine elongations that are not reflected in known measures like the circularity index.

**Figure S1:**
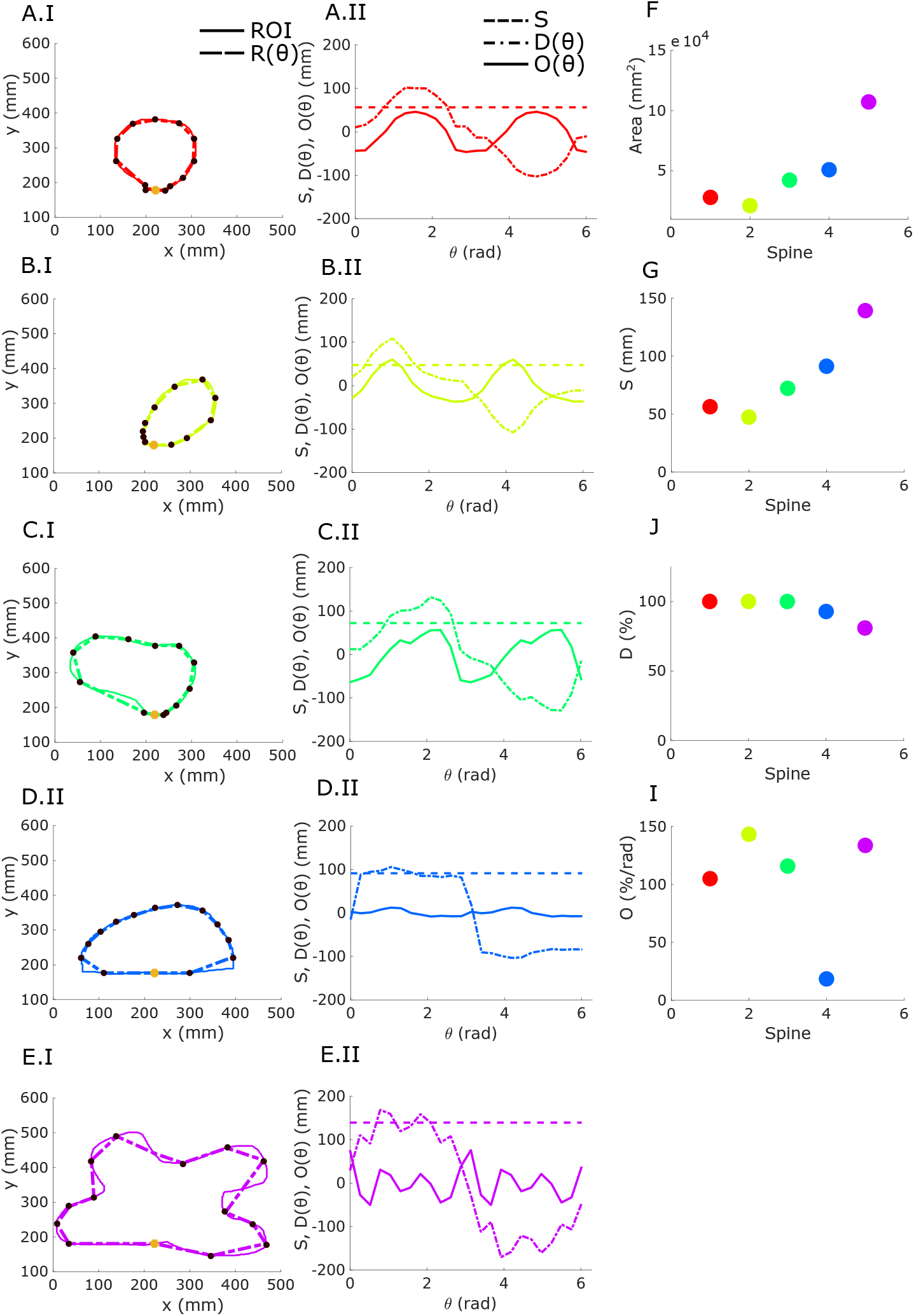
Shape descriptors. Left column: Different spine head shapes (solid line) and their approximation function *R*(*θ*) from Eq. (1) (dashed line). Black asteriks represent sample points and yellow dot the neck center. Center Column: Different parts of *R*(*θ*). Here the dashed, dotted dashed, and solid lines represent *S*, *D*(*θ*), and *O*(*θ*), respectively. Right column: Value of shape descriptors for the different spines (color-coded).

**Figure S2:**
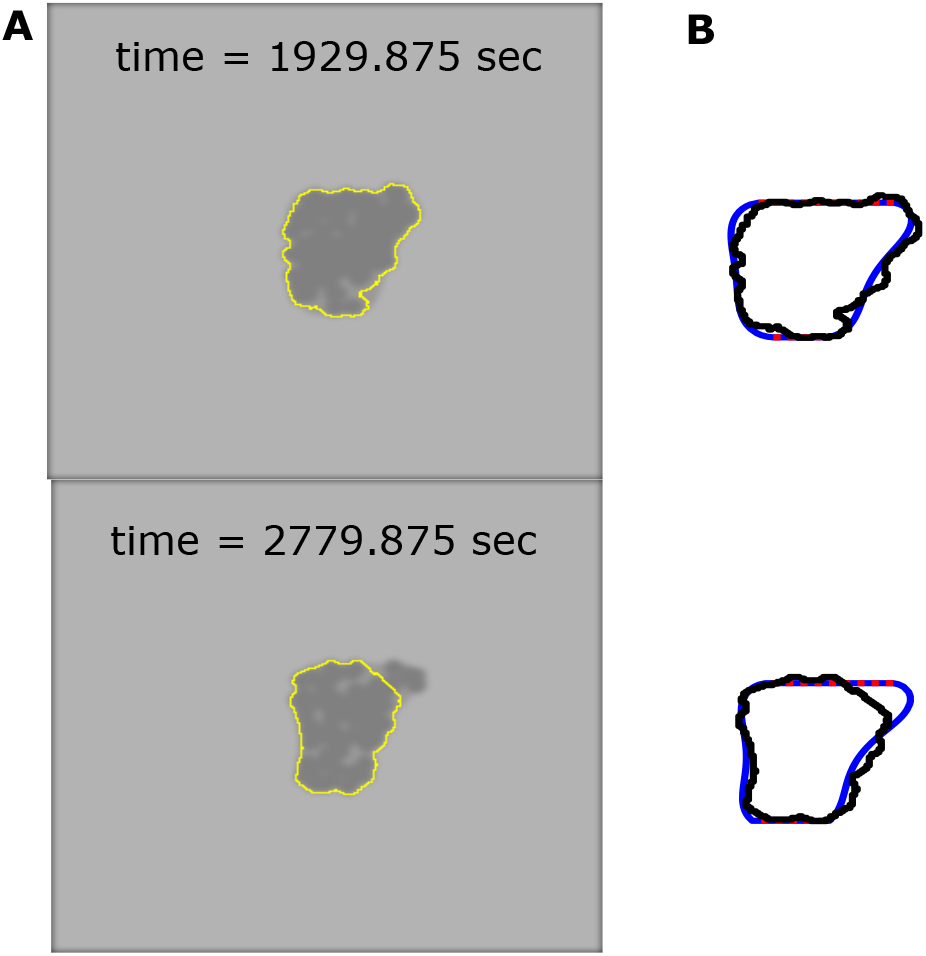
Model replicating imaging effects. **A** Model spine shape from the simulation in Figure 5 with fluorophores (circles) after filtering. Yellow line corresponds to the traced ROI following the methology described in Methods. **B** Extracted ROI from A and original shape. Red dotted lines correspond to the neck and PSD.

### Difference between model and experimental data

Experimental data suffers from some inaccuracies due to the imaging method used. For example, in confocal microscopy images reflect the florescence of certain particles that are not equally distributed and appear blurred, depending on the quality of the microscope. To replicate such effects in our modeled spines, we randomly allocate fluorophores inside the spine shapes, represented by small circles (see Fig. S2A) colored with a dark grey. The inside of the spine and the figure background are colored in different shades of grey to mimic experimental recordings. Then, the image is blurred using the `imgaussfilt` function in Matlab with *σ* = 1.99, which corresponds to the standard deviation of the point spread function calculated from the spines corresponding to the top stack in Figure 1B using Fiji. The resulting images are imported to Fiji and a ROI is traced, as in the experimental data. Next, the shape descriptors are calculated in Matlab (see values in Table S2). Note that the values from the blurred images change considerably from those of the model.

When analyzing all the blurred sample shapes of the simulation, we observe an increase in the coefficient of variance. However, the ROIs extracted from experimental data using the mRuby2 channel and the merged image show similar CVs, thus, the variance is conserved independently of the position of the fluorophores’ locations (see Fig. S3). Therefore, we assume that the variability difference between experimental data and simulations is due to inaccuracies in the imaging method.

**Table S2:**
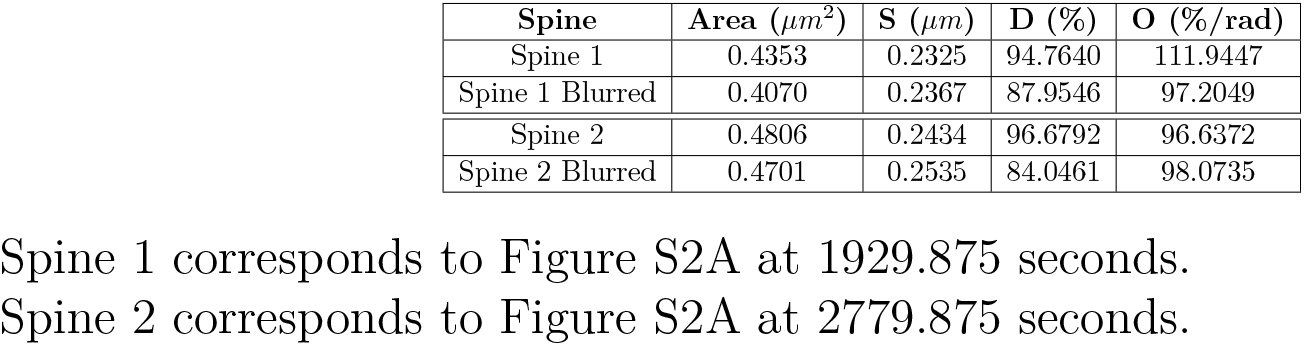
Shape Descriptors

| Spine | Area ($\mu m^2$) | S ($\mu m$) | D (%) | O (%/rad) |
| --- | --- | --- | --- | --- |
| Spine 1 | 0.4353 | 0.2325 | 94.7640 | 111.9447 |
| Spine 1 Blurred | 0.4070 | 0.2367 | 87.9546 | 97.2049 |
| Spine 2 | 0.4806 | 0.2434 | 96.6792 | 96.6372 |
| Spine 2 Blurred | 0.4701 | 0.2535 | 84.0461 | 98.0735 |

Spine 1 corresponds to Figure S2A at 1929.875 seconds.
Spine 2 corresponds to Figure S2A at 2779.875 seconds.

**Figure S3:**
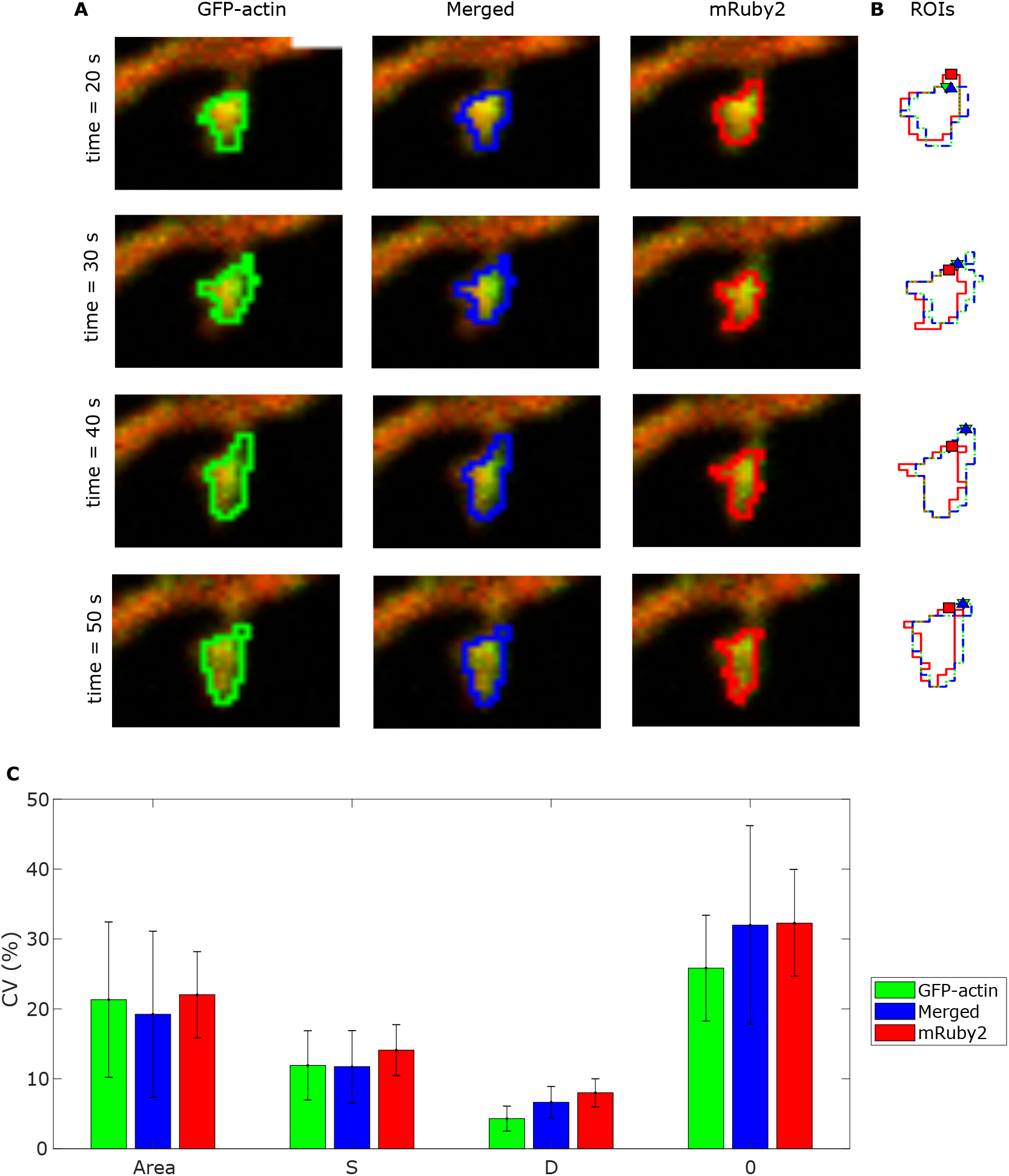
ROIs from experimental data. **A** ROIs extracted from experimental data usign the GFP-actin channel (right), as in Figure 5, mRuby2 channel (left) and both channels merged (center). **B** ROIs from A overlapped. **C** Median ± standard deviation of the coefficient of variability for shape descriptors calculated for each spine in the experimental data.

### Self-organized Criticality Validation

We test whether the model self-organizes to a critical state without parameter finetuning. For this, we vary the branching rate amplitude *ϕ* that linearly relates to the branching probability, given by Eq. (11) (see Methods). Hence, *ϕ* modifies the number of barbed ends. Figure S4 shows that the log-log plots of the probability of avalanche size, lifetime and avalanche interval show a power law behaviour. Note that the probability lifetime of the avalanches has a smaller exponent with the new set of parameters (slope of magenta line in Fig. S4C-D). This is expected since, on the one hand, a smaller value of *ϕ* decreases the branching probability, and hence, the polymerization foci last less. On the other hand, an increase in *ϕ* increases the branching probability; however, the branching probability is nonlinear and inversely proportional to the number of barbed ends. Hence, when the number of barbed ends increase, the branching probability and, consequently, the foci lifetime decreases. Therefore, under this parameter variation the duration of avalanches is shorter, and the probability of finding longer avalanches is smaller.

We performed additional criticality tests to the simulations, as in Tetzlaff et al. (2010). First, we test if the system is temporally scale-free by changing the size of the time bins. Figure S5A-C shows that the avalanche size in simulations with different parameters had power law distribution regardless of the time bin. To test whether the system is spatially scale-free, we assume that, due to experimental limitations, we observe only two or one foci at a time. Thus, the avalanche size is calculated only with those foci: if a third focus starts when two are active, the third one is discarded and the barbed ends of that focus are not included in the avalanche size calculation. The avalanche size distribution in Figure S5D-F exhibits a power law relation, hence, the system is spatially scale-free.

To evaluate if the interval between avalanches displays a scale-free behavior, two variables are introduced: *s_c_* the critical size of an avalanche and *η_c_* the waiting time between to avalanches with size bigger than *s_c_*. To compare the distributions of *η_c_* given *s_c_*, we need to re-scale *η_c_*, because larger avalanches are less probable. Thus, *η_c_ → η_c_R*(*s_c_*), with *R*(*s_c_*) the rate of having an avalanche bigger than *s_c_* per time unit, and *P* (*η_c_, s_c_*) *→ P* (*η_c_, s_c_*)*/R*(*s_c_*). Note that after re-scaling the distributions in Figure S5G-I collapse into a single function *F*, hence *P* (*η_c_, s_c_*) = *R*(*s_c_*)*F* (*η_c_s_c_*). Note that *F* is a power law, confirming that the intervals between avalanches are scale-free.

**Figure S4:**
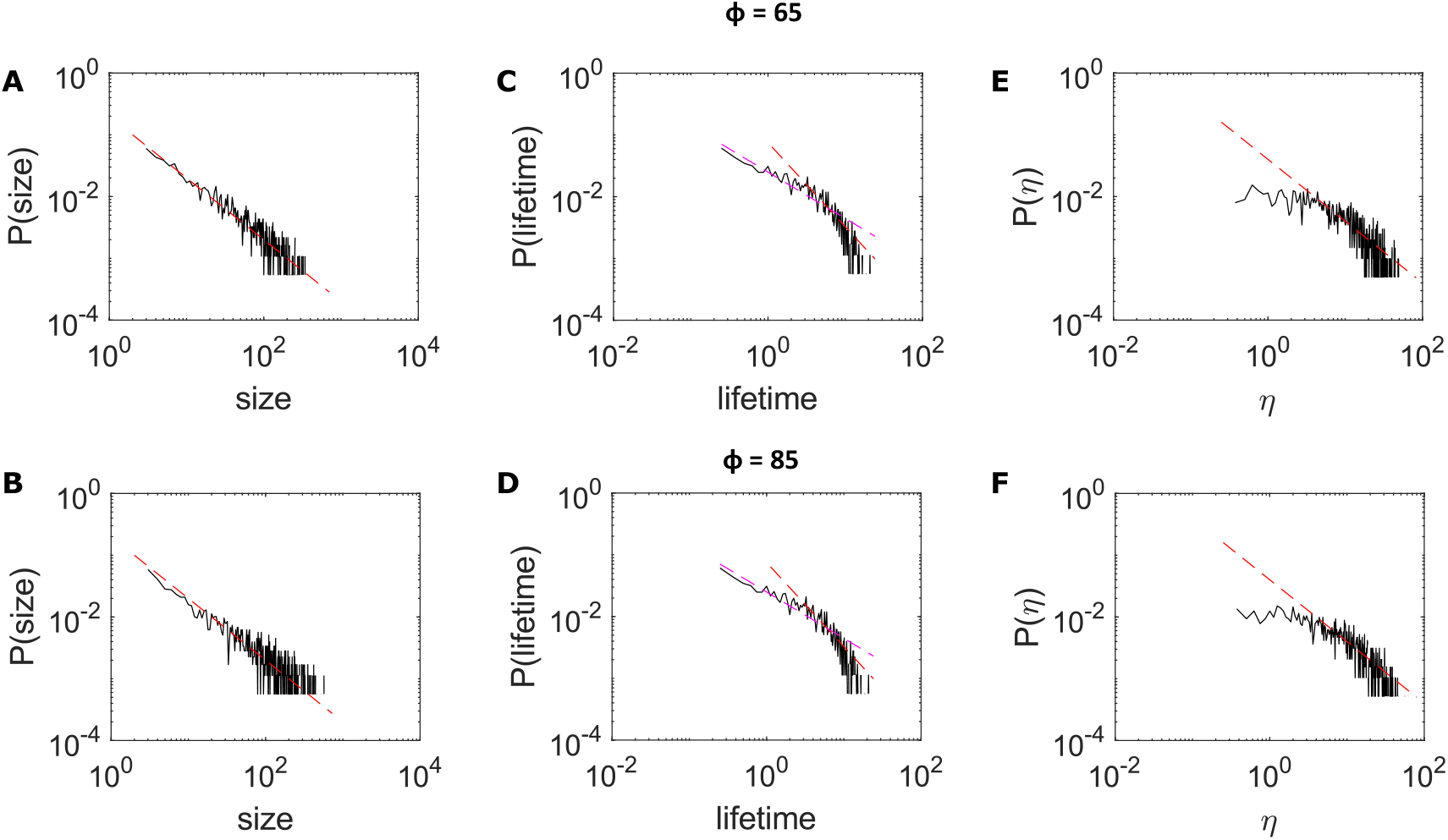
Self-organized criticality under parameter variation. Top *ϕ* = 65, bottom *ϕ* = 85. Log-log plot of the probability of avalanche size **A-B**, lifetime **C-D**, and duration of the interval between avalanches **E-F** corresponding to the data from 10 simulations of the model with the corresponding parameters. Red dotted lines correpond to straight lines with slope of −1 (A-B and E-F) and −1.36 (C-D). Magenta dotted lines have slope of −0.75.

Finally, we obtain the Allen Factor to assess criticality because it allows calculation of power law exponents *α* in the range of 0 *< α <* 3 (García-Marín et al., 2008). We define *S_z_*(*T*) as the number of polymerizing events contained in a time window *z* of length *T* and the Allen Factor

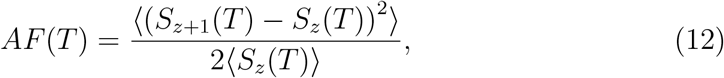

where ⟨·⟩ denotes the expectation value. In short, the Allen Factor measures the degree of clustering of the polymerization events compared to a homogeneous Poisson point process, in which *AF* (*T*) = 1, *∀T*. Note that in Figure S5J-K the Allen Factor has power law behavior for several time windows *T*, which indicates a scale-free point process.

**Figure S5:**
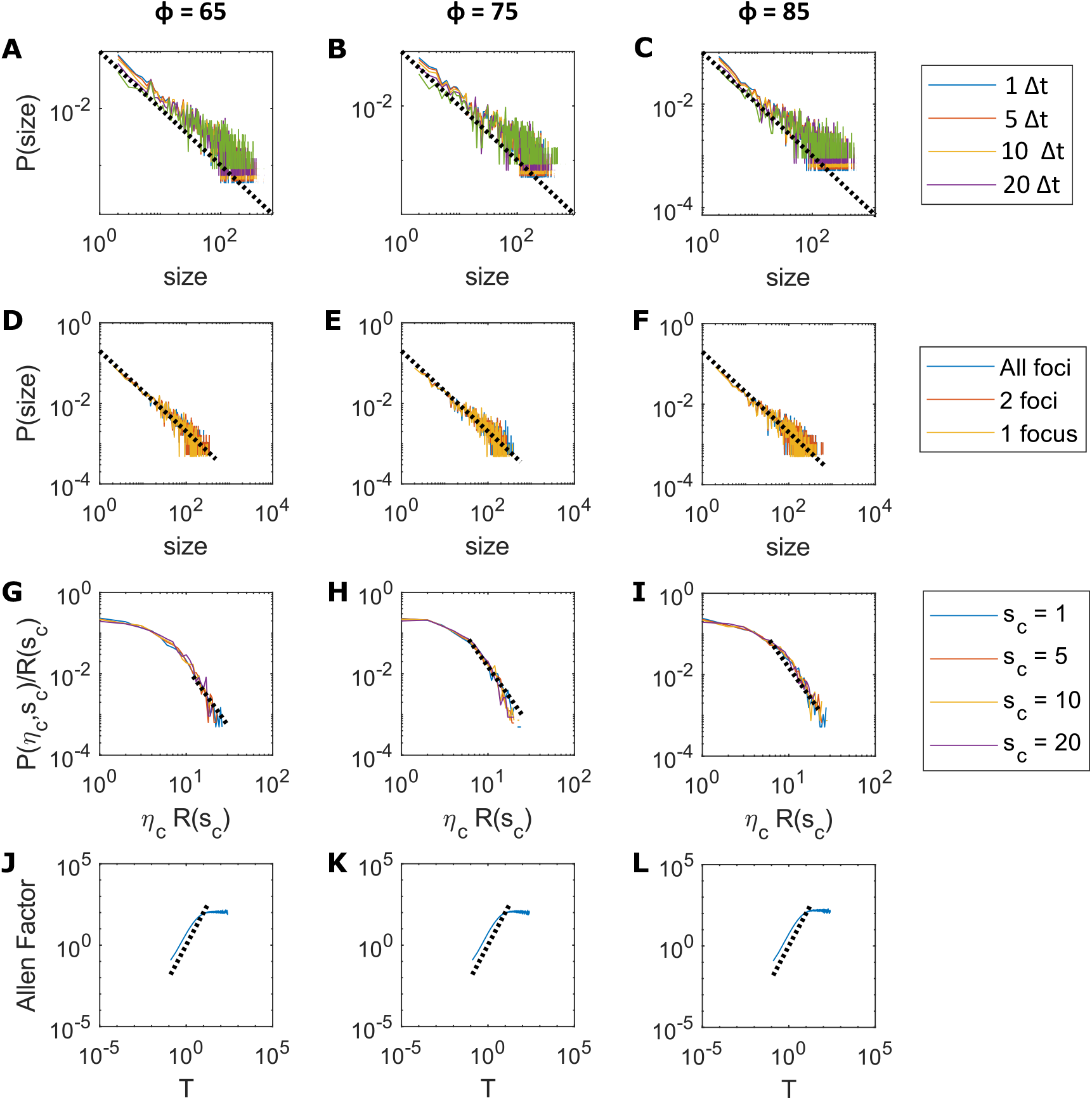
Self-organized criticality tests. Left, center, and right correspond to simulations with *ϕ* = 65, *ϕ* = 75, and *ϕ* = 85, respectively. Log-log plot of the probability of avalanche size taking different time bins **A-C** and restricting the number of foci **D-F** for the measurement. **G-I** Probability of interval duration *η_c_* between avalanches of size bigger than *s_c_* (see text for details). **J-L** Allen Factor for different time windows *T*. Black dotted lines correpond to straight lines with slope of −1 (A-F), −3 (G-I), and 2 (J-I). Data from 10 simulations of the model.

### Model Parameters

**Table S3:** Model Parameter Values.

| Symbol | Unit | Definition | Value |
| --- | --- | --- | --- |
| $\Delta_t$ | s | length of the time-step | 1/8 |
| $\delta_s$ | $\mu\text{m}$ | target length of and edge | 0.03 |
| $r_{neck_0}$ | $\mu\text{m}$ | initial neck radius | 0.0995 |
| $r_{PSD_0}$ | $\mu\text{m}$ | initial PSD radius | 0.3571 |
| $h_{neck_0}$ | $\mu\text{m}$ | initial value for fixing the neck | -0.49 |
| $h_{PSD_0}$ | $\mu\text{m}$ | initial value for fixing the PSD | 0.35 |
| $n_{f_0}$ | 1 | initial number of nucleation points | 4 |
| $\gamma_{cap}$ | $\text{s}^{-1}$ | barbed-end capping rate | 1 |
| $\gamma_{uncap}$ | $\text{s}^{-1}$ | uncapping rate for - ends | 1/30 |
| $\gamma_{sever}$ | $\text{s}^{-1}$ | depolymerisation/Severing rate of - ends | 1 |
| $\gamma_f$ | $\text{s}^{-1}$ | nucleation rate of new actin focus of activity | 0.1 |
| $a$ | $\mu\text{M}$ | concentration of profilin-ATP-actin at steady state | 3.8 |
| $\phi$ | $\mu\text{m}^{-2}$ | proportionality constant | 75 |
| $k_{on}$ | $\mu\text{M}^{-1}\text{s}^{-1}$ | barbed-end monomer assembly rate constant | 11.6 |
| $\delta$ | $\mu\text{m}$ | length of an actin monomer | 0.0022 |
| $k_B T$ | $\text{pN}\mu\text{m}$ | thermal energy | 0.0041 |
| $P$ | $\text{pN}\mu\text{m}^{-2}$ | difference between internal and external pressure | 85.7143 |
| $\tau$ | $\text{pN}\mu\text{m}^{-1}$ | surface tension | 15 |
| $\kappa$ | $\text{pN}\mu\text{m}$ | bending modulus | 0.18 |
| $\alpha$ | $\text{pN}$ | strength of filament influence | 3.8 |
| $\sigma$ | 1 | extend of filament influence | 0.2 |
| $\zeta$ | $\mu\text{m}^2\text{s}^{-1}\text{pN}^{-1}$ | strength of force update | 0.002 |
| $\lambda$ | $\mu\text{m}$ | nucleation distance parameter | 0.025 |

Table Modified from (Bonilla-Quintana et al., 2020).

